# Role of High Mobility Group B protein HmbA, orthologue of yeast Nhp6p in *Aspergillus nidulans*

**DOI:** 10.1101/2022.05.04.490617

**Authors:** J. Ámon, G. Varga, I. Pfeiffer, Z. Karácsony, Z. Hegedűs, C. Vágvölgyi, Z. Hamari

## Abstract

The mammalian HMGB1 protein belongs to the high-mobility-group B (HMG-B) family, which is not only architectural but also functional element of the chromatin. The fungal counterpart of HMGB1 was identified in *Saccharomyces cerevisiae* as Nhp6p and the pleiotropic physiological functions of this protein were thoroughly studied during the last decades. Although filamentous Ascomycete fungi also possess the orthologues of Nhp6p, their physiological functions, apart from their role in the sexual development, have not been investigated, yet. Here we study the physiological functions of the Nhp6p orthologue HmbA from *Aspergillus nidulans* in the primary and secondary metabolism, stress tolerance, hypha elongation and maintenance of polarized growth through the analysis of *hmbA* deletion mutant. We also revealed that the endochitinase ChiA acts in the cell wall remodelling and contributes to polar growth. Additionally, by conducting heterologous expression studies, we further demonstrated that HmbA and Nhp6p is interchangeable for several functions. We hypothesized that the fully complemented functions might predominantly depend on the DNA binding ability of the HmbA and Nhp6p proteins rather than on the interaction of these HMG-B proteins with other functional protein components of the chromatin.

## Introduction

Chromatin structuring high-mobility-group B (HMG-B) proteins are not only architectural but also functional elements of the chromatin (reviewed in (Ueda and Yoshida, 2010)). Architectural HMG-B proteins typically harbour two or more copies of HMG-box domains that establish interaction with DNA by a non-sequence-specific manner and also with various protein components of the chromatin. They are not only architectural components of the chromatin but they facilitate the establishment of stabile protein-DNA interaction between chromatin associated activator or repressor proteins with their cognate DNA motif, henceforth they greatly contribute to the normal functioning of the chromatin (involving transcription, repression, repair, recombination etc., reviewed in (Reeves, 2010)).

The first studied architectural HMG-B proteins in fungi were the non-conventional homologues of mammalian HMGB1 (earlier HMG1/2), the Nhp6Ap and Nhp6Bp paralogues from *Saccharomyces cerevisiae* (Kolodrubetz and Burgum, 1990; Kolodrubetz et al., 1988). Unusually, these two paralogue yeast proteins have only one copy of HMG-box domain and they possess a relatively short, basic N-terminal tail. The basic N-terminal region establishes electrostatic interactions with the DNA phosphodiester backbone of the major groove and the L-shaped HMG-box domain composed of three helices binds into the minor groove through both electrostatic and hydrophobic interactions (Allain et al., 1999; Yen et al., 1998). These protein-DNA interactions result in a sharp bend on the DNA backbone or stabilize the already distorted or non-B-type DNA (Allain et al., 1999; Paull et al., 1996). These interactions with the DNA together with the establishment of protein-protein interactions with other chromatin-associated proteins enable the Nhp6 proteins to promote the assembly of nucleosomes, DNA-protein complex formation of various transcription factors, activators, remodelling complexes or repressors (Celona et al., 2011; Hargreaves and Crabtree, 2011; Hepp et al., 2017; Laser et al., 2000; Paull et al., 1996). Nhp6 proteins participate in activation of transcriptions by RNA polymerase II and activation of *SNR6* by RNA polymerase III (Dowell et al., 2010; Kruppa et al., 2001; Lopez et al., 2001; Paull et al., 1996). Nhp6 has two functionally redundant paralogues, the Nhp6Ap and Nhp6Bp proteins; therefore the physiological role of the Nhp6 proteins could be assessed through the study of *NHP6A/NHP6B* double deletion mutants (*nhp6AΔBΔ*) (Costigan et al., 1994). The physiological effect of the lack of Nhp6 proteins resembles to that observed in *Slt2Δ* mutant and it was shown that *NHP6* is a target of the Slt2/Mpk1 MAPK signal transduction route. The *nhp6AΔBΔ* strain is reduced in growth, sensitive to N-starvation, sensitive to 38 °C that is suppressed by supplementing the medium with 1 M sorbitol and shows various morphological aberrations, such as elongated buds and enlarged bud-necks with enhanced chitin deposition. Transcriptome analyses of *NHP6^+^* and *nhp6AΔBΔ* strains revealed a pleiotropic effect for the lack of the Nhp6 proteins as several hundreds of genes display an altered expression in the *nhp6AΔBΔ* strain (Dowell et al., 2010; Moreira and Holmberg, 2000).

Orthologous counterpart of the yeast Nhp6p from *Aspergillus nidulans* is the HmbA protein (Karacsony et al., 2014). It was shown recently that HmbA plays a pivotal role in the sexual development and the proper expression of the mating-type genes (Bokor et al., 2019). Cleistothecia are formed in *hmbAΔ* however they are devoid of ascospores (Bokor et al., 2019). Although the *hmbA* gene deletion mutant and the deletion reconstituted strain were obtained in our previous work, which revealed the essential role of HmbA in sexual reproduction (Bokor et al., 2019), the overall physiological function of HmbA is still unexplored. Here we thoroughly examined the physiological functions of HmbA through the analysis of primary and secondary metabolism, stress sensitivity and micromorphology-related phenotypes in the *hmbA* deletion mutant and investigated the functional interchangeability of HmbA and Nhp6 by heterologous expression of *hmbA* and *NHP6A* genes in yeast *nhp6AΔBΔ* and *A. nidulans hmbAΔ* mutants, respectively.

## Results and Discussion

### HmbA is structurally similar to Nhp6Ap

Despite an NMR-based model structure of the Nhp6Ap-DNA complex is available (Allain et al., 1999), we aimed to create a ligand-free Nhp6Ap structure model together with the HmbA structure for comparative purposes (Table 1). A large fraction (67.74%) of the AAs (amino acids) is identical between HmbA and Nhp6Ap, and the three dimensional (3D) structures of the proteins also considerably overlap, especially at the three helices (H1-H3) (table 1 and figure 1). The H1 and H2 helices are highly similar between HmbA and Nhp6Ap (13 and 12 AAs are identical out of the 16 and 14 AA long helices, respectively), while the H3 helices are less similar (10 AAs out of the total 29 AAs of Nhp6A H3 helix are identical in HmbA) (figure 1). Notably, none of the altering AA residues of the helices map to the DNA binding interface; all such AAs map to the outer surface of the molecule as it is demonstrated in figure 1. These AA differences between HmbA and Nhp6A result in different surface electrostatic topography, which might be a deterministic property for the establishment of interaction with certain protein components of the chromatin (figure 1). Absolute conservation of the DNA-binding interface in HmbA compared to Nhp6Ap includes the i) R19-K22 (RKKK) region, responsible for binding the major groove (R13-K16 in Nhp6Ap); ii) P27 (P21 in Nhp6Ap) that directs the N-terminal tail toward the major groove (Yen et al., 1998); iii) S32 and Y34 (S26 and Y28 in Nhp6Ap) that establish H-bonds with the bases of the minor groove and is supposed to be responsible for a certain level of sequence specific DNA binding (Allain et al., 1999); and iv) M35 and F54 (M29 and F48 in Nhp6Ap) that intercalate into the major groove (Allain et al., 1999; Yen et al., 1998). According to the above data, the DNA binding properties of the yeast and *A. nidulans* HMG-B proteins are supposedly similar, while their capability to interact with other protein components of the functional chromatin is supposedly different.

**Figure 1.**
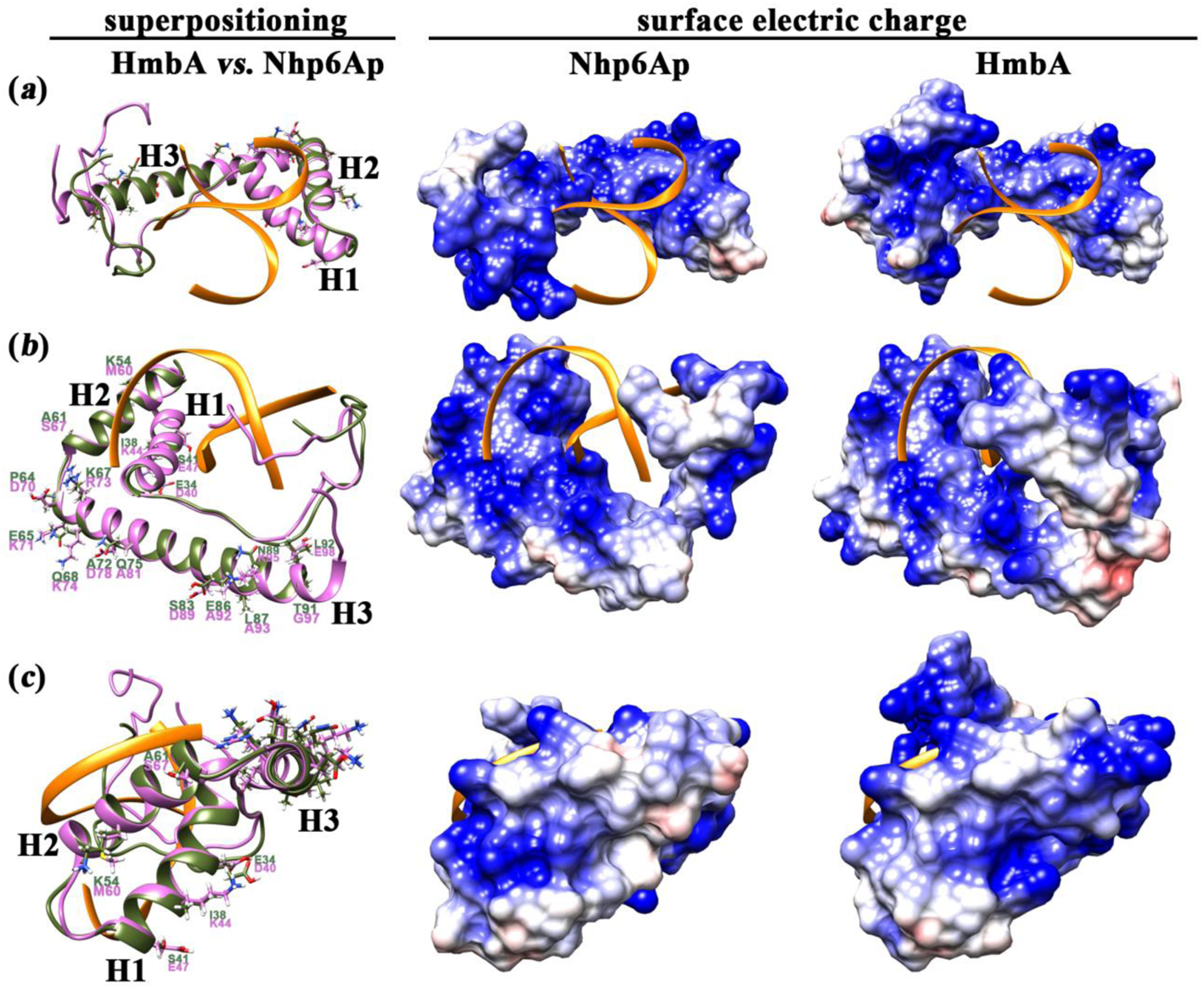
Comparison of the structure and electrostatic surface topography between HmbA and Nhp6Ap. The left side of each panel show the superpositioning of HmbA (pink ribbon) with Nhp6A (olive ribbon) from different angles, whereas the middle and right side of each panel correspond to Nhp6Ap and HmbA, respectively and show the electrostatic potential of the exposed molecule parts. AA residues that differ between the helices of HmbA and Nhp6A are indicated by showing their side chains in stick view. (*a*) The DNA binding interface is shown. None of the altering AA residues map to the DNA binding interface. (*b*) H2 and H3 helices are shown as well as the N-termini and the negatively charged C-terminus that is characteristic only to HmbA. (*c*) H1 and H2 helices are shown. H3 helix is shown from a top view, which illuminates that the altering AA residues are positioned to the surface of the molecule. Yellow ribbon depicts the DNA ligand. H1, H2 and H3 denote helices from N to C termini. Electrostatic potential from −10 (red colour) to +10 (blue colour) of the modelled proteins is shown. White colour indicates uncharged area.

**Table 1.** Results of HmbA and Nhp6A protein modelling and superpositioning of these models

| Quality assessment of the protein models | HmbA (106 AAs) | Nhp6A (93 AAs) |
| --- | --- | --- |
| C-score of initial model | 1.31 | 0.28 |
| estimated TM-score of initial model | 0.55±0.15 | 0.68±0.12 |
| estimated RMSD of initial model | 6.8±4.0 Å | 4.4±2.9 Å |
| TM-score of refined model to initiate model | 0.9881 | 0.9935 |
| RMSD of refined model to initiate model | 0.446 Å | 0.285 Å |
| RAMA<sup>1</sup>, % of AAs in favoured region | 93.3%<br>(84 AAs) | 96.3%<br>(78 AAs) |
| RAMA<sup>1</sup>, % of AAs in allowed region | 4.4%<br>(4 AAs) | 1.2%<br>(1 AA) |
| RAMA<sup>1</sup>, % of AAs in disallowed region | 2.2%<br>(2 AAs) | 2.5%<br>(2 AAs) |
| number of non-Pro, non-Gly residues | 90 AAs | 81 AAs |
| **Quality assessment of the superpositioning of the modelled HmbA and Nhp6Ap proteins (by built-in Matchmaker of UCSF Chimera 1.14)** |  |  |
| Sequence alignment scores<sup>2</sup> | 361.5 |  |
| Structure RMSDs between 75 pruned atom pairs in superposition | 0.509 Å |  |
| Structure RMSDs between all 93 atom pairs in superposition | 4.725 Å |  |
| Structure RMSDs across all 89 fully populated columns in the final alignment | 1.248 Å (SDM (cutoff 5.0): 24.766; Q-score: 0.685) |  |
| modelled interacting molecule in the PDB model | DNA |  |
| Overall identity between two proteins | 67.74% |  |

<sup>1</sup> RAMA: Ramachandran plot analysis results on non-Proline and non-Glycine regions derived from using Procheck server (https://servicesn.mbi.ucla.edu/PROCHECK/)
<sup>2</sup> Computing secondary structure assignments of superimposed models used ksdssp (Kabsch and Sander Define Secondary Structure of Proteins) with the following parameter values: -0.5 energy cutoff; minimum helix length 3; minimum strand length 3. Sequence alignment scores were obtained by Matchmaker (built in UCSF Chimera 1.14) with the following parameter values: chain pairing: bb; alignment algorithm: Needleman-Wunsch using BLOSUM-62 matrix; ss (secondary structure) fraction: 0.3; gap opening penalties (HH/SS/other)(HH: intra Helix; SS: intra Strand) 18/18/6, gap extension penalty: 1; ss scoring matrix: (O, S): -6 (H, O): -6 (H, H): 6 (S, S): 6 (H, S): -9 (O, O): 4 (H is Helix, S is Strand, O is Other); iteration cutoff: 2.

However, we might not exclude the possibility of having common interacting protein partners between HmbA and Nhp6Ap if the interaction depends on the surface AAs of the most conserved H1 and H2 helices (figure 1*b,c*). The yeast and *A. nidulans* proteins have a rather striking difference at the C-terminal end. The HmbA protein ends in a negatively charged unstructured tail that might have a special functional role, while Nhp6Ap lacks such unstructured tail (its carboxy terminal practically ends with the H3 helix) (figure 1*b*).

Proper expression of *SNR6* gene in yeast depends on Nhp6p proteins, which expression was found to be vital at 38 °C (Kruppa et al., 2001; Lopez et al., 2001). Interaction of Nhp6Ap with the promoter *SNR6* facilitates the binding of TFIIIC to its suboptimally spaced *SNR6* promoter elements (Kruppa et al., 2001). While the absence of Nhp6Ap is tolerated at 30 °C (TFIIIC can bind, although with a reduced capacity, to the *SNR6* promoter elements), the binding of TFIIIC to the suboptimally spaced *SNR6* promoter elements at 38 °C is not supported without Nhp6p (Kruppa et al., 2001). The *SNR6* gene expression is downregulated in an *nhp6AΔBΔ* strain at 30 °C, which is ceased to zero at 38 °C (Kruppa et al., 2001). The *nhp6AΔBΔ* mutant shows a reduced growth at 30 °C, whereas such a mutant is non-viable at 38 °C (Costigan et al., 1994; Kruppa et al., 2001; Lopez et al., 2001). The heterologous expression of the *A. nidulans hmbA* from the native *NHP6A* promoter in the *nhp6AΔBΔ* strain (yC’*hmbA*) significantly improved the expression of *SNR6*, and cured the heat sensitive phenotype (figure 2*a,b*). Success of the restoration of *SNR6* regulation and growth at 38 °C by HmbA confirms that the DNA binding ability of HmbA is not different from that of Nhp6Ap.

**Figure 2.**
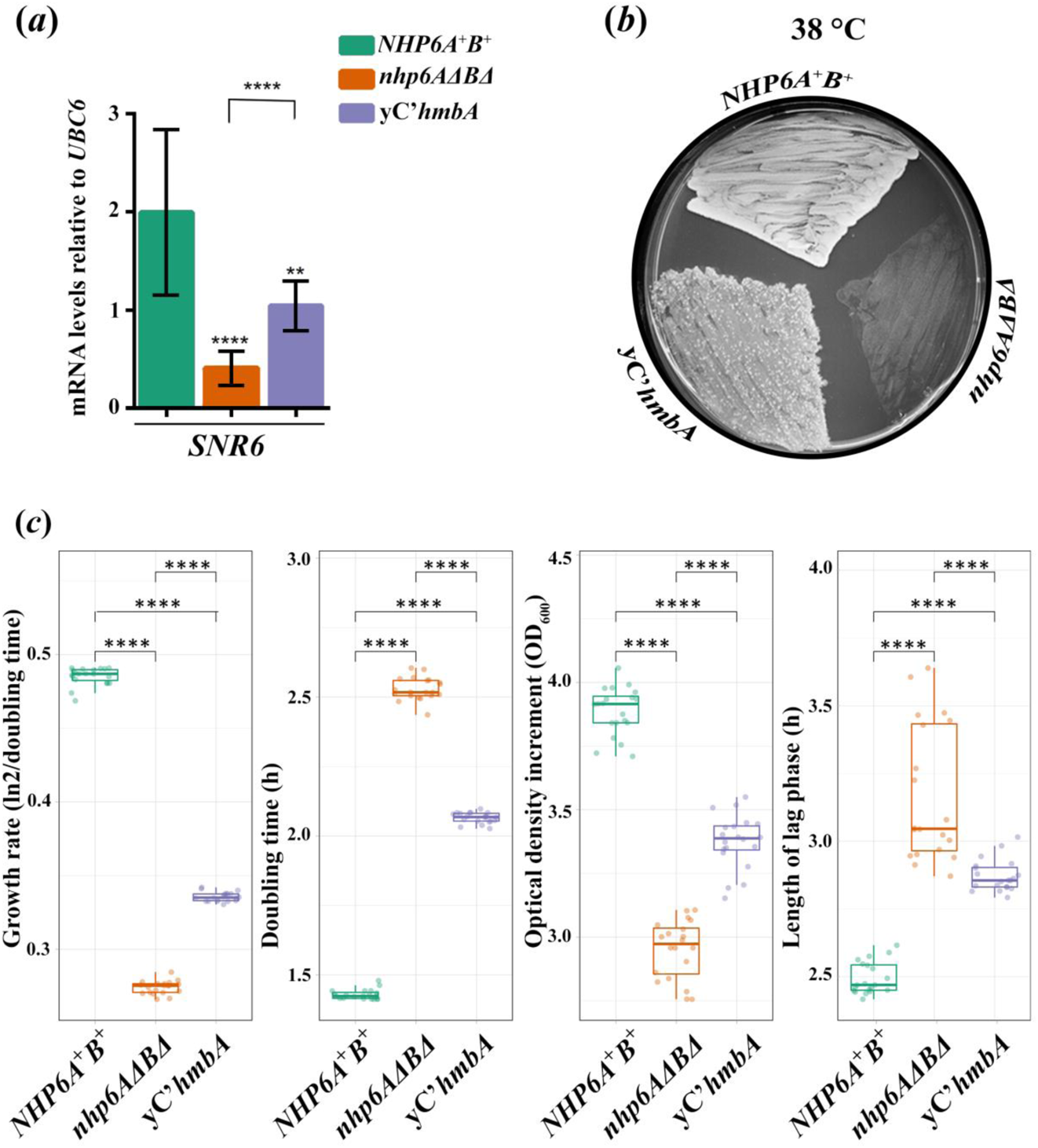
Downregulation of *SNR6*, heat sensitivity and reduced fitness of yeast *nhp6AΔBΔ* strain is remediated by the heterologous expression of *A. nidulans hmbA*. (*a*) mRNA levels of *SNR6* in *NHP6A^+^B^+^*control (Y199), *nhp6AΔBΔ* (HZS.891) and yC’*hmbA* (*nhp6AΔBΔ* complemented with *hmbA*, HZS.890) strains measured by RT-qPCR. Strains were grown on YPD up to OD_600_=0.3-1.0 at 38 °C. RT-qPCR data were processed according to the standard curve method (Larionov et al., 2005) with *UBC6* as the reference mRNA. The mean and standard deviations of three independent experiments are shown. Significance between strains was assessed by using Student’s *t*-test. **/**** indicate *P* < 0.01/0.0001. Complete genotypes of the strains and used primers are listed in electronic supplementary material, table S1 and S2, respectively. (*b*) Heat sensitive phenotype of *nhp6AΔBΔ* is remediated by the heterologous expression of *A. nidulans hmbA*. Strains (same as described in panel *a*) were grown at 38 °C on YPD. (*c*) Results of fitness measurements including growth rate, doubling time, optical density increment, and length of the lag phase. Used strains are described in panel *a*. Significance between strains was assessed by using Mann-Whitney U-test. **** indicates *P* < 0.0001.

Remediation of further *nhp6AΔBΔ* phenotypes, such as prolonged doubling time was also investigated in yC’*hmbA* strain. We found that doubling time and additional fitness characteristics such as growth rate, optical density increment and length of lag phase are also remediated in the *hmbA* expressing *nhp6AΔBΔ* strain (figure 2*c*). Although the complementation was not complete, the yC’*hmbA* strain showed significant improvement in all tested fitness component (p<0.0001).

### Heterologous expression of *NHP6A* restores ascospore production in fruiting bodies of *hmbAΔ*

We constructed an *NHP6A* complemented *hmbAΔ* strain (C’*NHP6A*) where *NHP6A*, the yeast counterpart of *hmbA*, is expressed from the *hmbA* promoter (P*_hmbA_*) (for details see the Materials and methods). Since cleistothecia of the *hmbAΔ* strain are devoid of ascospores (Bokor et al., 2019), content of crushed C’*NHP6A* cleistothecia was monitored under microscope, which revealed that the ascospore formation was completely restored upon the heterologous expression of *NHP6A*. The same restoration of deletion phenotype was reported by us earlier in an *hmbA* complemented *hmbAΔ* strain (C’*hmbA*), where *hmbA* was expressed under the control of its physiological promoter (P*_hmbA_*) (Bokor et al., 2019).

### *hmbA* is steadily expressed from germination to mycelial growth and the encoded protein localizes in the nucleus

Germination (up to 3-4 h of development) is accompanied by the steady increase in *hmbA* mRNA level and the expression is steadily high throughout mycelial growth (5 and 6 h) (figure 1*a*). HmbA was localized in the nucleus based on co-localization with the DAPI stained nucleus (figure 1*b*).

### HmbA has influence on primary metabolism and response to environmental stresses

Deletion of *hmbA* results in reduced growth on complete medium due to the decrease in the elongation rate of the hyphae (Bokor et al., 2019). To test the effect of *hmbA* deletion on the primary metabolism, utilization of various carbon- and nitrogen-sources was compared between *hmbAΔ* and *hmbA^+^*control and C’*hmbA-gfp* (*hmbAΔ* complemented with *hmbA^+^-gfp*), C’*hmbA^+^* (*hmbAΔ* complemented with *hmbA^+^*) and C’*NHP6A* (*hmbAΔ* complemented with yeast *NHP6A*) reconstituted strains (figure 4). The growth tests were performed using glucose-nitrate medium at 37 °C as a control condition. On this medium *hmbAΔ* showed a significantly reduced growth (*P* < 0.0001, Mann-Whitney U-test) compared to both the *hmbA^+^* and all types of complemented strains that restored the *hmbA^+^*-like growth (electronic supplementary material, figure S1*a*). Response of all examined strains to the tested conditions was calculated by strainwise normalization of the colony sizes to the control condition (that is indicated in the electronic supplementary material, data). To assess significant differences in the response of *hmbAΔ* and in the various reconstituted strains to that of the *hmbA^+^* control we used Student’s *t*-test and Mann-Whitney U-test. In figure 4 we present both the growth tests and the comparative analysis of the responses to various selected conditions (for all tested conditions see electronic supplementary material, figure S1 and data). Using nitrate as the sole nitrogen-source, utilization of 9 various carbon-sources was compared to that of glucose (electronic supplementary material, figure S1). Utilization efficiency of sucrose, lactose, galactose and ethanol as sole carbon-sources was significantly decreased (*P* < 0.05-0.001) in *hmbAΔ* compared to *hmbA^+^* control (Figure 3*a*, electronic supplementary material, figure S1*b*).

**Figure 3.**
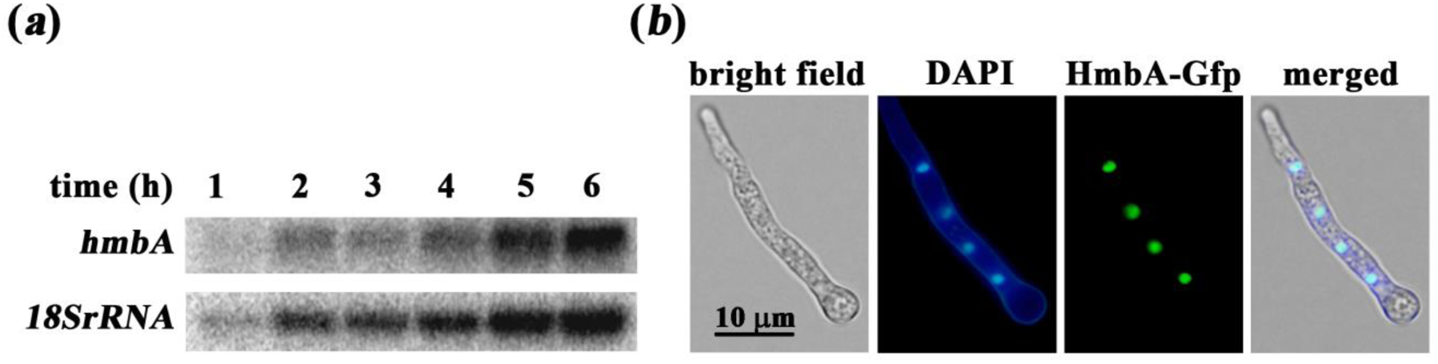
Expression profile of *hmbA* and intracellular localization of the gene product. (*a*) Northern blot analysis of the mRNA level of *hmbA* from germination to mycelial growth (1-6 h). Loading of RNA was estimated by hybridization with an 18S rRNA specific DNA probe. (*b*) Subcellular localization of HmbA-Gfp with DNA (DAPI) staining of young hypha in strain HZS.371. Conidiospores were germinated for 8 h in MM at 37 °C. Cells were stained for DNA (DAPI) and examined by fluorescence microscopy (Zeiss 49 and 15 filter sets were used for DAPI and GFP, respectively). Scale bar represents 10 μm.

**Figure 4.**
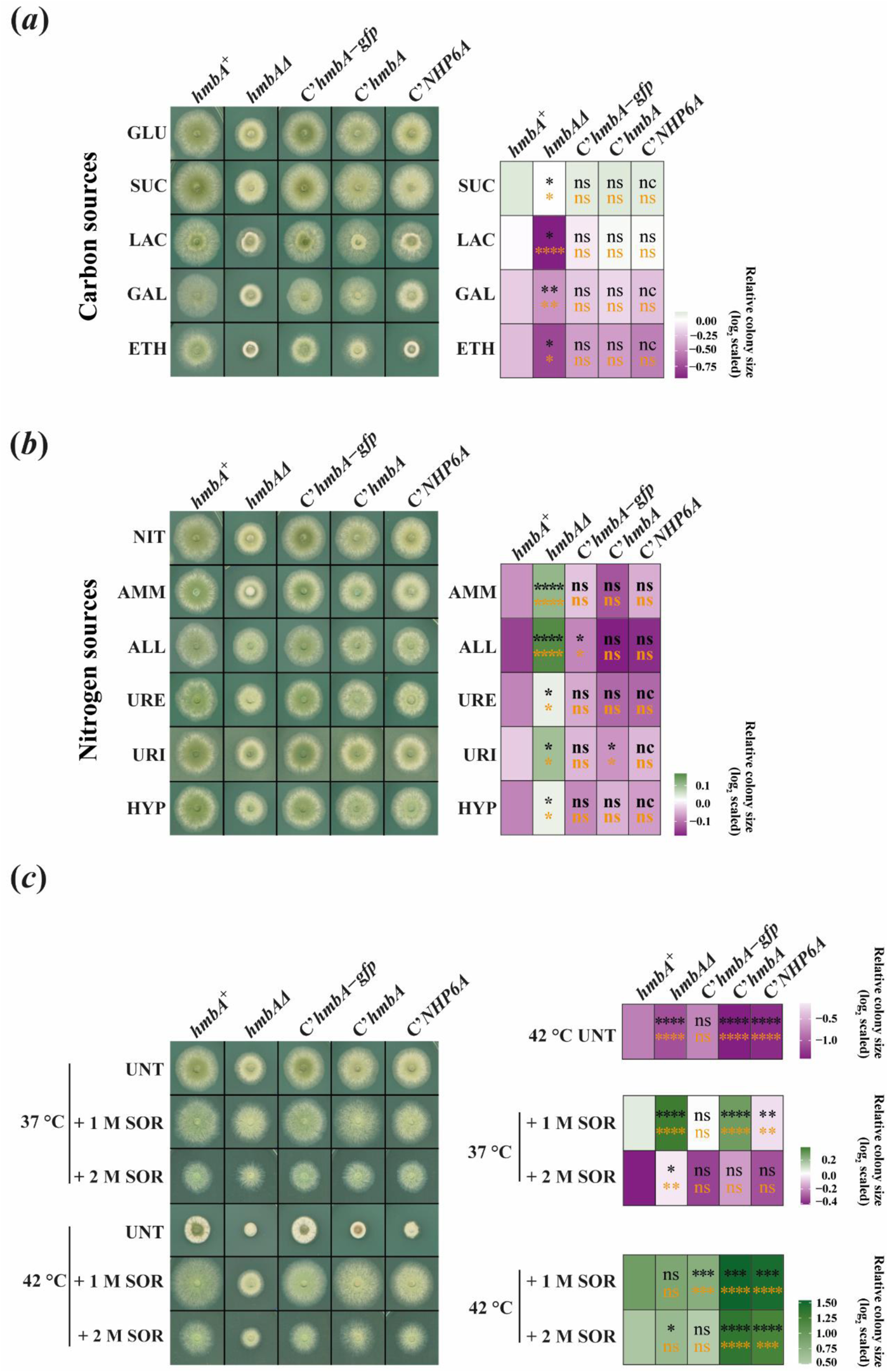
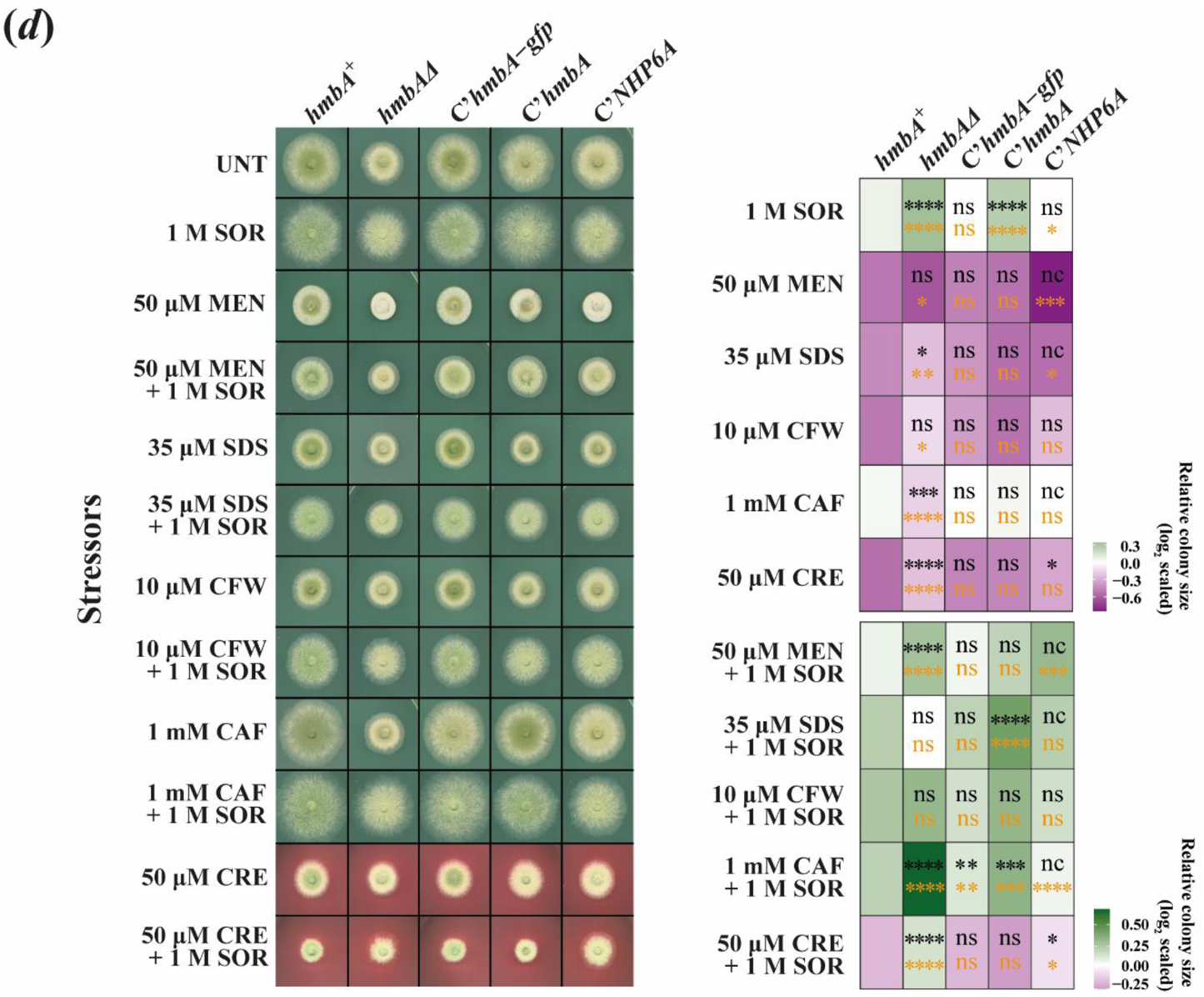
Growth tests of *hmbA^+^*, *hmbAΔ* and complemented control strains on various carbon- and nitrogen-sources and under various environmental stress conditions. On the right side of each growth tests, a heat map indicates the response of a strain to the tested conditions (log_2_(colony size_tested condition_/size_control condition_)). Asterisks in the cells show the significance of Student’s *t*-test (orange) and Mann-Whitney U-test (black) calculated by comparing the responses of all tested strains to that of the *hmbA^+^* control (*/**/***/**** indicates *P* < 0.05, 0.01, 0.001 and 0.0001, respectively; nc: non-calculated due to low sample size, ns: non-significant). (*a*) Carbon-source utilization tests on GLU (glucose), SUC (sucrose), LAC (lactose), GAL (galactose) and ETH (ethanol). The nitrogen-source was sodium nitrate in all conditions of panel *a*. (*b*) Nitrogen-source utilization test on NIT (sodium-nitrate), AMM (diammonium L-(+)-tartrate), ALL (allantoin), URE (urea), URI (uric acid) and HYP (hypoxanthine) as sole nitrogen-sources. The carbon-source was glucose in all conditions of panel *b*. (*c*) Effect of heat stress at 42 °C on growth of the tested strains without (UNT, untreated) or in the presence of 1 M sorbitol (1 M SOR) as osmotic stabilizer or 2 M sorbitol (2 M SOR) as osmotic stressor. All medium used in panel *c* was a glucose/sodium nitrate MM (minimal medium). (*d*) Growth test on glucose-sodium nitrate MM (UNT, untreated) supplemented with various stress agents in the indicated concentrations without or in the presence of 1 M sorbitol (1 M SOR) osmotic stabilizer. MEN, menadione; SDS, sodium dodecyl sulphate; CFW, Calcofluor White; CAF, caffeine; CR, congo red. Used strains were *hmbA^+^* as control (HZS.120); *hmbAΔ* (*hmbA* deletion strain, HZS.320); *C’hmbA* (*hmbAΔ* complemented with *hmbA*, HZS.621), C’*hmbA-gfp* (*hmbAΔ* complemented with *hmbA-gfp*, HZS.371); C’*NHP6A* (*hmbAΔ* complemented with *NHP6A*, HZS.834). Genotypes of the strains are listed in the electronic supplementary table S1.

Deletion of *hmbA* did not affect the utilization of glycine maltose, raffinose, sorbitol and xylose (electronic supplementary material, figure S1*b*). Using glucose as the sole carbon-source, utilization of 6 various nitrogen-sources was compared to that of nitrate (Figure 3*b*, electronic supplementary material, figure S1*c*). Utilization of diammonium L-(+)-tartrate, allantoin, urea, uric acid and hypoxanthine as sole nitrogen-sources by *hmbAΔ* was significantly better (*P* < 0.05-0.0001) compared to that by *hmbA^+^*control. Utilization of acetamide as sole nitrogen-source was not significantly different between *hmbA^+^* and *hmbAΔ*. Utilization of the tested carbon- and nitrogen-sources in the complemented strains was not significantly different from the *hmbA^+^*control, except that the C’*hmbA-gfp* strain utilized allantoin better (*P* < 0.05) compared to the *hmbA^+^* control whereas C’*hmbA^+^*utilized uric acid as sole nitrogen-source less efficiently (*P* < 0.05) than the *hmbA^+^* control did.

Severe heat stress (42 °C) affected the *hmbAΔ* growth more adversely than that of the *hmbA^+^* strain (*P* < 0.0001) (figure 4*c*). Although most of the *hmbA* deletion phenotype is restored by complementing the deletion strain with *hmbA* or yeast *NHP6A* (see primary metabolism- and secondary metabolism-related phenotypes), the heat sensitivity was not restored to the wild-type-like level in C’*hmbA^+^*and C’*NHP6A* strains. Notably, complementation of *hmbAΔ* by *hmbA-gfp* (C’*hmbA-gfp*) restored the heat sensitivity to the wild-type level (Figure 3*c*, electronic supplementary material, figure S1*d*). We reasoned that this restoration stem from having two copies of the *in trans*-integrated *hmbA-gfp* construct, which provided a better support against the heat-stress compared to the support the *hmbA^+^* or *NHP6A* construct did in single copy. Supplementation of the medium with 1 M sorbitol remediated the heat sensitivity at 42 °C in all tested strains (Figure 3*c*, electronic supplementary material, figure S1*d*). While sorbitol acts as osmotic stabilizer when used in 1 M concentration, doubling the concentration (2 M sorbitol) causes an osmotic stress. At 37 °C, the *hmbAΔ* tolerated the osmotic stress on 2 M sorbitol better (*P* < 0.01) than the *hmbA^+^* or the complemented controls did.

Oxidative stress agent menadione in 50 µM concentration was less tolerated by *hmbAΔ* compared to *hmbA^+^*(*P* < 0.05) (figure 4*d*, electronic supplementary material, figure S1*e*). Cell wall disruptors (10 µM Calcofluor White, 50 µM Congo Red, 35 µM SDS) were better tolerated by *hmbAΔ* compared to *hmbA^+^* (*P* < 0.05, *P* < 0.0001 and *P* < 0.01, respectively), however *hmbAΔ* was more sensitive to 1 mM caffeine compared to *hmbA^+^* (*P* < 0.0001) (figure 4*d*, electronic supplementary material, figure S1*e*). The differential responses of *hmbAΔ* and *hmbA^+^* in the presence of cell wall disruptors might reflect strain-specific differences in the cell wall architecture. The effect of 1 M sorbitol on growth remediation was similar in all tested strains (figure 4*d*, electronic supplementary material, figure S1*e*). Sensitivity of *hmbAΔ* strain against heavy metal stress (50 µM cadmium sulphate) was similar to that of *hmbA^+^* and the C’*hmbA* and C’*hmbA-gfp* reconstituted strains (electronic supplementary material, figure S1*e*). Notably, the C’*NHP6A* reconstituted strain showed extreme sensitivity to cadmium sulphate, which was rescued by the supplementation of the medium with 1 M sorbitol osmotic stabilizer (electronic supplementary material, figure S1*e*).

### Deletion of *hmbA* results in altered chitin deposition at the hyphal tips, production of thin hyphae and an abnormal mode of hypha elongation

Deletion of *hmbA* resulted in a decreased hyphal elongation rate on complete medium (Bokor et al., 2019) as well as on glucose-nitrate medium (figure 4*a*, electronic supplementary material, figure S1*a*). The slow growth is accompanied by a compact mycelium formation, composed from thin and aberrantly running (zig-zag shaped) hyphae (figure 5*a,e*). The hyphae of *hmbAΔ* frequently run in a markedly wavy course and have more side-branches compared to the *hmbA^+^*control (figure 5*a*). The C’*hmbA-gfp* and C’*hmbA* reconstitution strains displayed a restoration of the wild-type morphology, while the recovery was only partial in the C’*NHP6A* reconstitution strain (electronic supplementary material, figure S2). Since the growth tests on media supplemented with cell wall disruptor agents indicated that the cell wall architecture might be different in *hmbAΔ* from that of *hmbA^+^* control (figure 4*d*), the distribution of chitin at the hyphal tips was monitored by Calcofluor White staining. Chitin staining was intense at the top edge of the wild type hyphae that was gradually decreased in the sub-apical area (figure 5*b*). Such a wild-type like diffuse chitin distribution in the sub-apical area was rarely observed in the *hmbAΔ* hypha tips, which indicates that remodelling of cell wall at the subtending zone below the apex is damaged in *hmbAΔ*. Remodelling of the cell wall, especially at the subtending zone, requires the action of endochitinases and endoglucanases (Martinez-Nunez and Riquelme, 2015). We found that *chiA* (codes for endochitinase supposedly involved in cell wall remodelling, (Takaya et al., 1998)) is significantly downregulated in the *hmbAΔ* mutant by 2.77 fold (*P* < 0.01) compared to the *hmbA^+^* control (figure 5*c*). To test whether *chiA* downregulation contributes to the aberrant hyphal growth phenotype, we overexpressed the *chiA* gene in an *hmbAΔ* strain (*hmbAΔ* OE*chiA*). Overexpression of *chiA* re-established the hypha elongation rate to normal (the colony size become wild-type-like), improved the hypha morphology and restored the diffuse distribution of chitin in the sub-apical area supporting the notion that the normal expression of *chiA* might be critical for the above processes (figure 5*a,b*).

**Figure 5.**
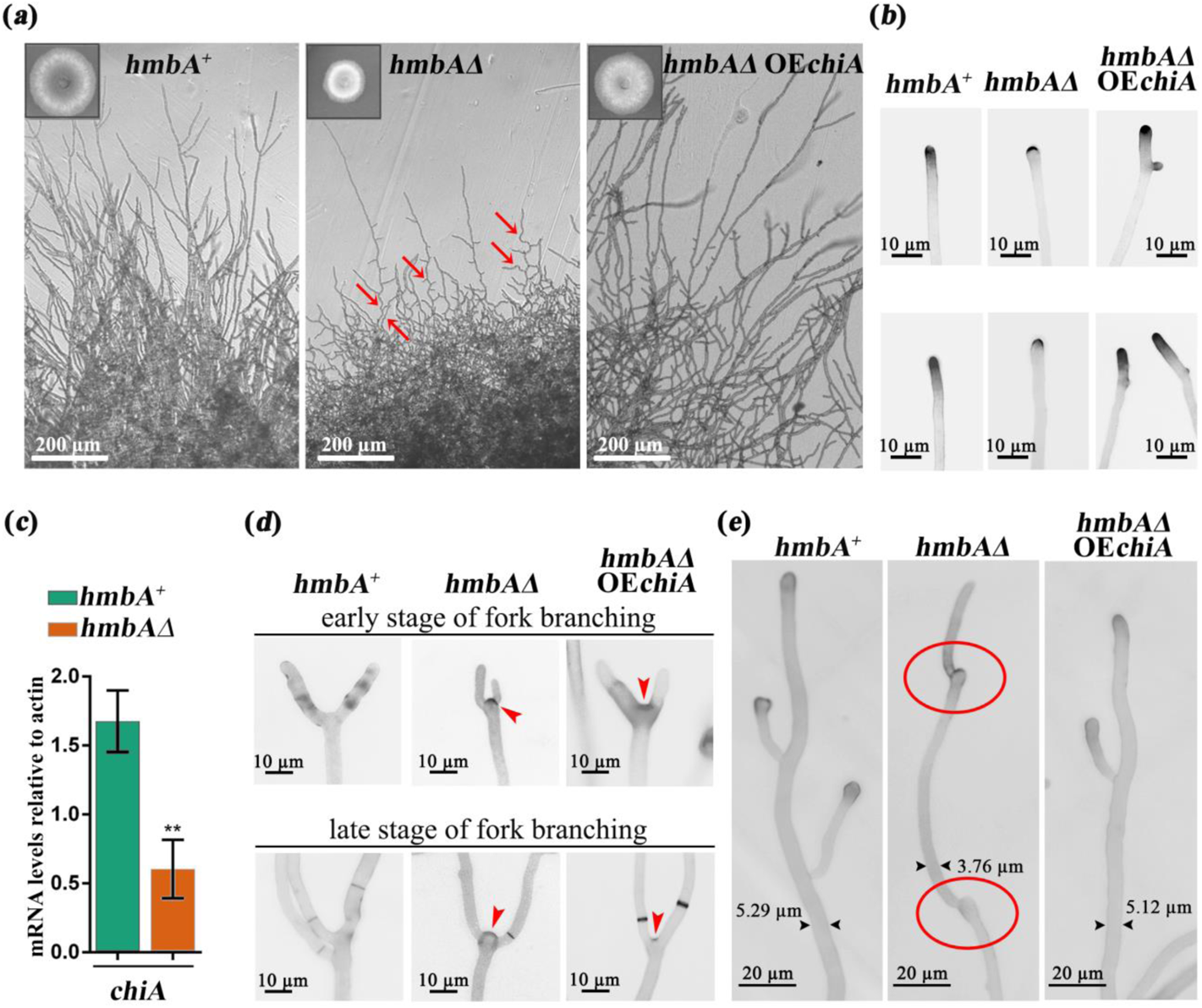
Downregulation of *chiA* altered morphology and chitin distribution in *hmbAΔ*. (a) The images show the morphology of elongated hyphae across three tested strains. Bright field images were documented with Leica DMI 4000B, DFC295 detector. Scale bars are shown. Red arrows denote zig-zag shaped hyphae. Insets show colonies grown on glucose-nitrate minimal medium at 37 °C for 3 days. (*b*) The images show Calcofluor White staining of chitin at the hyphal tips across three tested strains. Scale bars are shown. Microscopy was carried out with Zeiss Axioobserver 7 Axiocam 503 mono detector with DAPI filter setting. (*c*) The figure shows the mRNA levels of *chiA* in the tested strains measured by RT-qPCR. Strains were grown on glucose-nitrate minimal medium at 37 °C for 9 hours. RT-qPCR data were processed according to the standard curve method (Larionov et al., 2005) with γ-actin transcript (*actA*/AN6542) as reference. Standard deviations of three independent experiments are shown. Primers are listed in electronic supplementary material, table S2. (*d*) The images show the fork-branching at the hyphal tips of the tested strains at early and later stages of branching. Arrowheads denote the relicts of hyphal tips that underwent fork-branching. Scale bars are shown. (*e*) The images show hypha elongation across three tested strains. Samples were stained with Calcofluor White. Scale bars are shown. Red circles in midsections of elongated hyphae mark the relicts of arrested hyphal tips that underwent a germination-like process to continue the polar growth. This non-conventional way of elongation resulted in zig-zag shaped hyphae (denoted by red arrows in panel *a*). Tested strains were *hmbA^+^*(*hmbA^+^* control, HZS.120), *hmbAΔ* (*hmbA* deletion strain, HZS.320) and *hmbAΔ* OE*chiA* (*chiA* overexpressing *hmbAΔ* strain, HZS.921). The complete genotypes are listed in electronic supplementary material, table S1.

Notably, the nascent branching sites at the tips (fork-branching sites) in *hmbAΔ* retained the hyphal tip morphology together with the deposited chitin at the apical edge (figure 5*d*). The fork branches emerged similarly as germ tubes emerge from conidiospores. This phenomenon suggests that the normal mode of polarized growth is deficient in *hmbAΔ* and the mutant applies an alternative mode for hyphal tip growth. Overexpression of *chiA* in *hmbAΔ* strain restored the normal mode of polarized growth, although a low level of abnormal chitin deposition at the fork-branching sites was still detected (figure 5*d*).

According to the results presented above, downregulation of cell wall hydrolase endochitinase *chiA* leads to incomplete remodelling of the *hmbAΔ* cell wall at the subtending zone below the apex, which cannot further support the continuous elongation of the hyphal tips. The polarized growth of the hyphae is frequently arrested as it is clearly demonstrated in figure 5*e*. The abnormal structures at the mid-sections of the presented *hmbAΔ* hypha correspond to the relicts of earlier arrested hyphal tips. Resuming the polarized growth at the arrested tips seems to be possible in *hmbAΔ* by a mechanism that resembles to germination. The germination-like growth is most probably initiated at the arrested tips where the cell wall is the most weakened. Repeated occurrence of consecutive arrests and not-conventional resumptions of polarized growth results in the formation of zig-zag shaped hyphae (figure 5*a*). Overexpression of *chiA* in *hmbAΔ* restored the continuous and conventional mode of hypha elongation (figure 5i)

### Germination of *hmbAΔ* conidiospores is delayed

Deletion of *hmbA* resulted in 0.5 h delay in germination of conidiospores (figure 6*c*). Conversion of the trehalose deposited in resting conidiospores to glycerol is necessary for the swelling of conidiospores during the isotropic growth phase of germination (d’Enfert et al., 1999). According to this, we hypothesized that the delay of germination of *hmbAΔ* conidiospores might be associated with impaired trehalose content or impaired conversion of trehalose to glycerol. By measuring of the trehalose and glycerol content by HPLC in the first 120 min of germination (isotropic growth phase), we revealed that despite of the trehalose content of *hmbAΔ* resting conidiospores was significantly lower (*P* < 0.05) compared to that of *hmbA^+^*, the trehalose in *hmbAΔ* was metabolised with the same rate as in *hmbA^+^* after 30 min of incubation (figure 6*a*). Remarkably, the lower initial trehalose content of *hmbAΔ* conidiospores did not limit the glycerol production, moreover, significantly more (*P* < 0.05) glycerol was produced in the *hmbAΔ* conidiospores (figure 6*a*). Beside trehalose, various polyols (such as mannitol) could also serve as a source of the high level of glycerol (conversion of mannitol to glycerol during germination was demonstrated for *Aspergillus niger* ((Witteveen and Visser, 1995) and reviewed in (Meena et al., 2015)).

**Figure 6.**
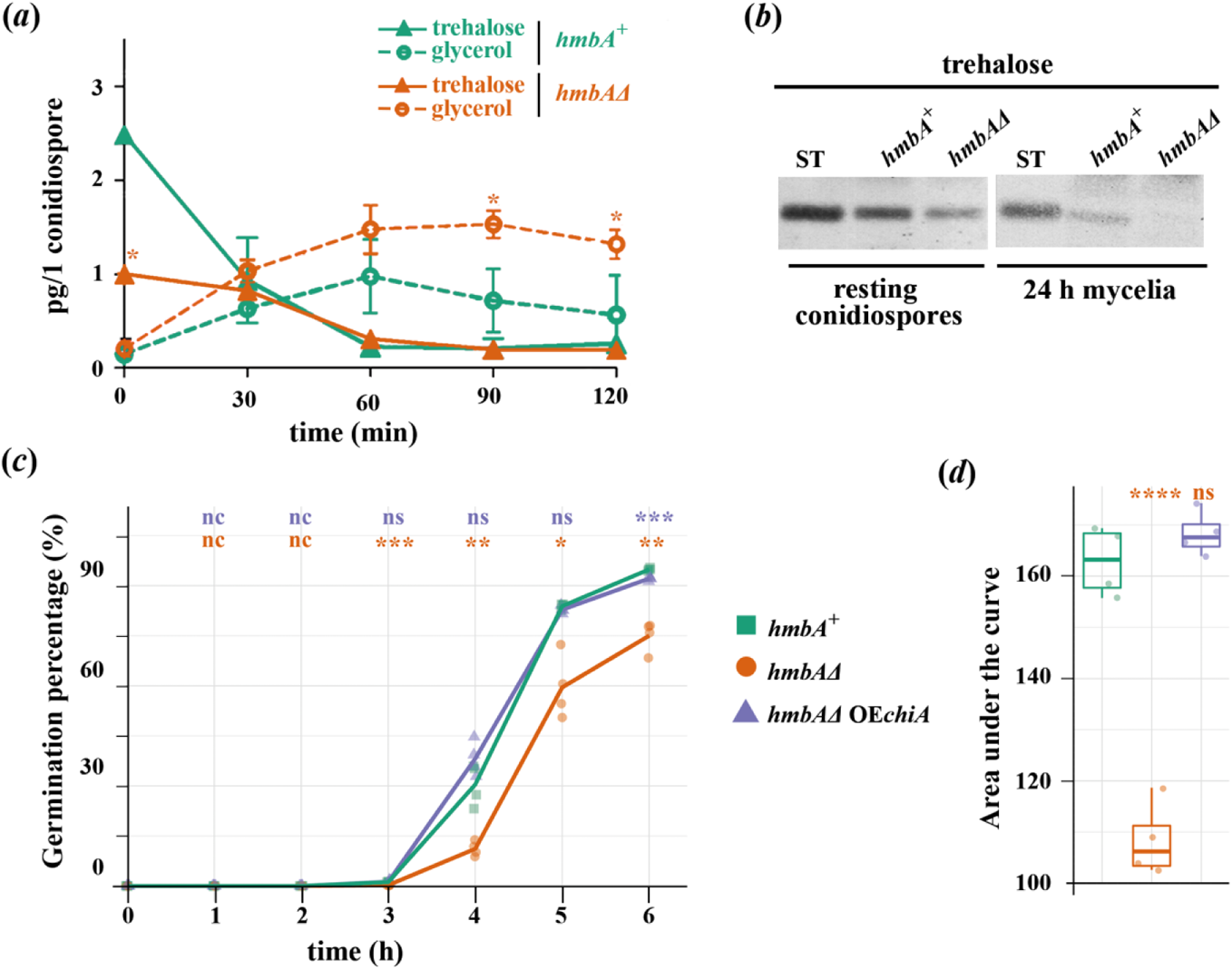
Comparison of trehalose content and metabolism of trehalose to glycerol in *hmbAΔ* and *hmbA^+^* conidiospores and characterization of conidiospore germination. (*a*) HPLC-detected conversion of trehalose to glycerol during the first two hours of germination in *hmbAΔ* and *hmbA^+^* conidiospores. Measurements started with the analysis of resting conidiospores at the 0 h time point. Colour and figure codes are shown in the figure legend. Mean of three biological replicates are shown. Error bars represent standard deviation. Significant differences (Student’s *t*-test) are marked with asterisks (* *P* < 0.05); (a) Detection of trehalose by TLC in conidiospores and mycelia of *hmbAΔ* and *hmbA^+^* strains. ST indicates trehalose standard. (*c*) Characterization of conidiospore germination rate in *hmbA^+^*, *hmbAΔ* and *hmbAΔ* OE*chiA* strains. 10^3^ conidiospores were inoculated in cellview cell culture dishes with glass bottom and incubated on 37 °C. Germination was monitored and documented by using Zeiss Axioobserver 7 microscope with Axiocam 503 mono detector. Germination percentage was calculated by normalizing the number of germinated conidiospores with the number of the non-germinated conidiospores. Solid lines denote the mean of four biological replicates. Colour and figure codes are shown in the figure legend. Significant differences (Student’s *t*-test) are marked with asterisks (* *P* < 0.05; ** *P* < 0.01; *** *P* < 0.001; nc: non-calculated due to each measured value is zero, ns: non-significant). Significance denoted by orange and blue correspond to pairwise comparison of the control *versus hmbAΔ* and control *versus hmbAΔ* OE*chiA* strain, respectively. (*d*) The figure shows the area under the curves presented in the germination rate graph (panel *c*) based on four biological replicates. Box plots show the median, first and third quartiles, with whiskers showing the 5^th^ and 95^th^ percentiles. Significant changes (Student’s *t*-test) are marked with asterisks **** *P* < 0.0001; ns: non-significant). Used strains were *hmbA^+^* as control (HZS.120); *hmbAΔ* (*hmbA* deletion strain; HZS.320) and *hmbAΔ* OE*chiA* (*chiA* overexpressing *hmbAΔ*; HZS.921). The complete genotypes are listed in electronic supplementary material, table S1.

Detection of trehalose content by TLC method further confirmed the decreased trehalose content of *hmbAΔ* conidiospores compared to that of *hmbA^+^* (figure 6*b*). Similar quantitative difference in trehalose content was also observed in the mycelia (figure 6*b*).

Since the level of glycerol is normal in the *hmbAΔ* conidiospores during the isotropic growth, the germination delay is not caused by any failure of swelling. After swelling, a certain area of the conidiospore cell wall must undergo a remodelling process where the germ tube can emerge and polar growth can begin. It is reasonable to assume that any impairment of cell wall remodelling might affect the germination. The endochitinase ChiA was reported to be required for normal germination by acting as a cell wall remodeller as deletion of *chiA* results in delayed germination of conidiospores and also for reduced growth of the *chiAΔ* colony (Takaya et al., 1998). This is in line with the impaired growth phenotype of *hmbAΔ* strain, in which the *chiA* expression is downregulated (figure 5*c*) and overexpression of *chiA* rescued the impaired growth phenotype of *hmbAΔ* (figure 5*a,b,d,e*) and concomitantly restored the germination dynamics to the *hmbA^+^*level (figure 6*c,d*).

### HmbA regulates the production of the secondary metabolite sterigmatocystin by a nitrogen source-dependent way

Sterigmatocystin (STC) production was barely detectable in *hmbAΔ* on complete medium and on lactose - ammonium minimal medium compared to the *hmbA^+^* control (figure 7). However, the produced STC was comparable with that in *hmbA^+^* on lactose - nitrate minimal medium (figure 7). Unravelling the nitrogen source-dependent role of HmbA in STC production is challenging, since multiple biological processes might be involved in (ranging from sensing and transducing of environmental signals to many aspects of interconnection of primary and secondary metabolism). Notably, even though *in trans* expression of *hmbA* and *hmbA-gfp* in *hmbAΔ* strain re-established the *hmbA^+^*-like STC production, the heterologous expression of *NHP6A* did not improve the STC production of *hmbAΔ* (figure 7).

**Figure 7.**
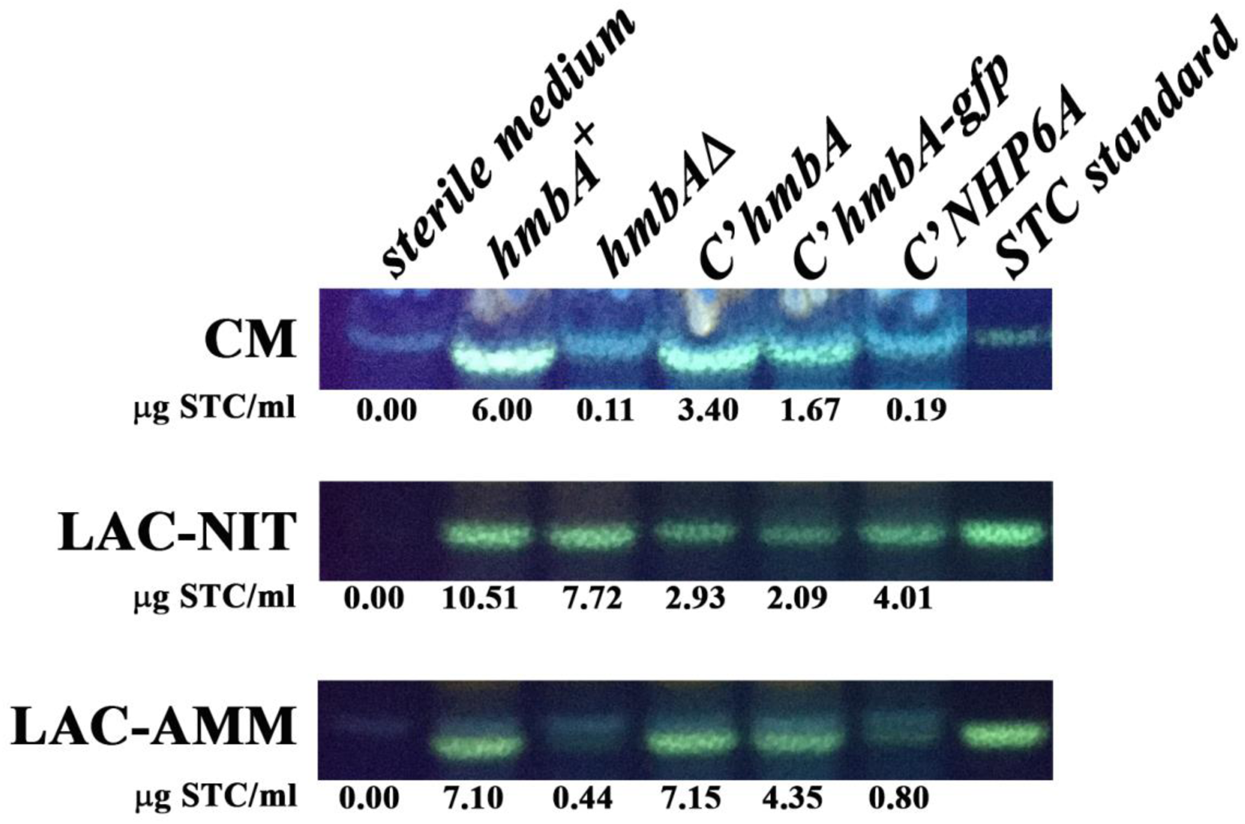
Sterigmatocystin production on various media in *hmbA^+^* control, *hmbAΔ* and various complemented *hmbAΔ* strains detected by TLC and quantified by HPLC. CM: complete medium, LAC-NIT: minimal medium with lactose carbon-source and sodium-nitrate nitrogen-source; LAC-AMM: minimal medium with lactose carbon-source and diammonium L-(+)-tartrate nitrogen-source. Used strains were *hmbA^+^*as control (HZS.120); *hmbAΔ* (*hmbA* deletion strain, HZS.320); *C’hmbA* (*hmbAΔ* complemented with *hmbA*, HZS.621), C’*hmbA-gfp* (*hmbAΔ* complemented with *hmbA-gfp*, HZS.371); C’*NHP6A* (*hmbAΔ* complemented with *NHP6A*, HZS.834). Sterile medium was used as negative control. STC is sterigmatocystin standard (Sigma). The complete genotypes are listed in electronic supplementary material, table S1.

## Conclusion

HmbA structurally corresponds to the HMGB1-like Nhp6p from *S. cerevisiae*. The DNA binding surface of HmbA is remarkably identical with that of Nhp6p, which makes the HmbA interchangeable with Nhp6A, at least in those functional roles (such as *SNR6* expression in yeast) where the role of the HMGB protein predominantly depends on its DNA-binding properties. Interchangeability of HmbA with Nhp6A was tested in many HmbA-affected physiological roles and in many cases, Nhp6A was able to complement the *hmbA* deletion phenotype (involving ascospore formation in the fruiting bodies, utilization of various carbon- and nitrogen-sources, radial growth rate, hypha elongation by mostly wild-type like polarized growth). We hypothesize that the physiological functions of HmbA, which are restored by the heterologous expression of *NHP6A* in an *hmbAΔ* strain predominantly depend on the DNA binding ability of the HmbA protein. However, there were physiological functions of HmbA, which were not complemented in a *hmbAΔ* strain by Nhp6Ap (including the response to menadione-generated oxidative stress and cadmium-sulphate generated heavy metal stress as well as the production of sterigmatocystin). This might reflect that these functions do not rely only on the DNA binding ability of HmbA, but also on protein-protein interactions between HmbA and other functional protein components of the chromatin (e.g., transcription factors).

We proposed a connection between the slow growth rate, the abnormal mode of polarized growth maintenance and the regulatory role of HmbA in the proper expression of cell wall remodellers, such as *chiA.* Apart from *chiA*, it is reasonable to assume that the expression of other cell wall remodeller genes are also influenced by HmbA and thereby the deficient polarized growth of *hmbAΔ* might be the net effect of the insufficient expression of multiple wall-remodellers. Earlier studies on the role of cell wall remodeller endochitinases in *A. nidulans* or in *A. fumigatus* failed to demonstrate the pivotal role of endochitinases in the maintenance of polar growth (Alcazar-Fuoli et al., 2011; Takaya et al., 1998; Yamazaki et al., 2008). The *hmbA* deletion phenotype provided a unique opportunity to reveal that cell wall remodelling endochitinases such as ChiA are pivotal for the cell wall remodelling and normal mode of maintenance of polar growth.

## Materials and methods

### Strains, cultures, growth conditions

The *A. nidulans* and *S. cerevisiae* strains used in this study are listed in the electronic supplementary material, table S1. Standard *Aspergillus* genetic markers, complete medium (CM) and minimal medium (MM) are described at the following URL: http://www.fgsc.net/Aspergillus/gene_list/. Media were supplemented with vitamins (www.fgsc.net) according to the requirements of each auxotrophic strain. The various carbon- and nitrogen-sources used in the experiments were the following: 1% (m/V) glucose (GLU), 1% (m/V) sucrose (SUC), 1% (m/V) galactose (GAL), 1% (m/V) xylose (XYL), 1% (m/V) maltose (MAL), 1% (m/V) lactose (LAC), 1% (m/V) sorbitol (SOR), 3.5% (m/V) raffinose (RAF), 2% (V/V) ethanol (ETH) and 2% (V/V) glycerol (GLY), 10 mM sodium-nitrate (NIT), 5 mM diammonium L-(+)-tartrate (AMM), 1 mM acetamide (ACE), 1 mM allantoin (ALL), 5 mM urea (URE), 0.6 mM uric acid (URI) and 1 mM hypoxanthine (HYP). Media with cell wall stressors and oxidative stress agents were supplemented in the following concentrations: 10 µM Calcofluor White (CFW), 50 µM congo red (CR), 35 µM sodium dodecyl sulphate (SDS), 50 µM menadione (MEN), 1 mM caffeine (CAF), 50 µM cadmium-sulphate (CDS). When indicated, media were supplemented with 1 M sorbitol as osmotic stabilizer. *S. cerevisiae* was maintained and studied on YPD medium (0.5% (m/V) yeast extract, 1% (m/V) peptone, 1% (m/V) glucose). During the genetic cross of yeast strains enriched sporulation medium (1% (m/V) potassium acetate (Fisher), 0.1% (m/V) yeast extract, 0.05% (m/V) glucose, 0.01% (m/V) amino-acids supplement powder mixture for sporulation) and MAT-a haploid selection medium (synthetic dropout medium: 0.5% (m/V) ammonium sulphate, 0.17% (m/V) Yeast Nitrogen Base, supplemented with 0.2% (m/V) glucose and with amino-acid mix, without histidine, arginine, and lysine) supplemented with 50 µg/ml canavanine (Sigma) and 50 µg/ml thialysine (Sigma) were used.

### Construction of *S. cerevisiae nhp6AΔBΔ* double deletion mutant

To generate a double deletion strain for *NHP6A* and *NHP6B*, first we obtained the *nhp6aΔ* (*nhp6aΔ::KanMX, his3Δ1 leu2Δ0 met15Δ0 ura3Δ0, MAT-a*) strain from the YKO Mat-a collection (Giaever et al., 2002) and the *nhp6bΔ* strain (*nhp6bΔ::NatMX, can1Δ::P_STE2_-SpHis5 lyp1Δ his3Δ1 leu2Δ0 ura3Δ0 met15Δ0, MAT-α*) from the SGA (Synthetic Genetic Array) query collection (Tong and Boone, 2006). To be able to use an expression construct at a later stage, we first swapped the KanMX deletion cassette of the *nhp6aΔ* strain to HphMX marker as follows. We performed a HindIII-EcoRI digestion of the pCRII-TOPO::hphMX plasmid and then after heat-inactivation, we transformed the plasmid into the *nhp6aΔ strain* using the standard lithium acetate method (Gietz and Schiestl, 2007). Transformants were selected on YPD supplemented with hygromycin (PAA Laboratories) at a final concentration of 400 µg/ml. The loss of the KanMX marker was confirmed by testing the inability of the transformants to grow on YPD supplemented with G418 (Sigma) at a final concentration of 200 µg/ml. We followed an established protocol (Tong and Boone, 2006) to generate the double mutant *nhp6AΔBΔ* by genetic cross. After several steps of selection, we used hygromycin and nourseothricin supplemented (400 µg/ml and 100 µg/ml, respectively) haploid selection medium to isolate MAT-a haploid double deletion mutants (*nhp6aΔ*::HphMX, *nhp6bΔ*::NatMX). Gene deletion was checked by “NHP6A confA frw”-“KanB rev” and “NHP6B confB frw”-“KanB rev” primers. Primers are listed in electronic supplementary material, table S2.

### Construction of *hmbA* expressing *nhp6AΔBΔ* yeast strain

In order to obtain the *hmbA* expressing *nhp6AΔBΔ* strain (yC’*hmbA*), the 321 bp cDNA of *A. nidulans hmbA* gene was ampified by “c-hmbA BamHI frw” and “c-hmbA SalI rev” primers and the BamHI-SalI digested PCR product was cloned into a BamHI-SalI digested yeast expression vector, M4801 (http://www.addgene.org/51664/). The P*_GAL_* promoter of the obtained vector (M4801-P*_GAL_*-C’*hmbA*) (electronic supplementary material, figure S3*a*) was truncated by AgeI-BamHI digestion and the 308 bp long promoter sequence of *NHP6A* (P*_NHP6A_*) was amplified (using “NHP6A prom AgeI frw” and “NHP6A prom BamHI rev” primers) and cloned into the AgeI-BamHI digested M4801-P*_GAL_*-C’*hmbA* vector. The resulted vector (M4801-P*_NHP6A_*-C’*hmbA*) (electronic supplementary material, figure S3*a*) was transformed into *nhp6AΔBΔ* strain (HZS.891) after NotI digestion using the standard lithium acetate method (Gietz and Schiestl, 2007). Correct transformants where the expression plasmid was integrated into the HO locus were selected on YPD supplemented with hygromycin, nourseothricin and G418 at a final concentration of 400 µg/ml, 200 µg/ml and 200 µg/ml, respectively. Integration of the vector was checked by “NHP6A prom AgeI frw”-“c-hmbA SalI rev”, while the copy number of integration events were checked with quantitative PCR according to Herrera et al. (Herrera et al., 2009) using “hmbA ReTi frw”-“hmbA ReTi rev” and the single copy control “UBC6 ReTi frw”-“UBC6 ReTi rev” primer pairs. Primers are listed in electronic supplementary material, table S2.

### Construction of *NHP6A, hmbA-gfp* and *chiA* expressing *hmbAΔ* strains

Transformation cassettes were constructed by using double-join PCR (DJ-PCR) method (Yu et al., 2004) and cloning. In order to obtain *NHP6A* complemented *hmbAΔ* strain (C’*NHP6A*), *NHP6A* gene was amplified from *S. cerevisiae* genome by using “nhp6A NcoI frw” and “nhp6A BamHI rev” primer pair (282 bp). The PCR product was digested with NcoI-BamHI and cloned into NcoI-BamHI digested pAN-HZS-1 vector (Karacsony et al., 2014) (the digestion eliminated the *gfp* gene from the vector). The obtained vector (pAN-HZS-18) expressed the *NHP6A* from the constitutive promoter P*_gpdA_*. In order to express *NHP6A* at the physiological level similar to that of *hmbA*, the P*_gpdA_*. promoter was truncated by NheI-NcoI digestion in pAN-HZS-18 that was followed by the cloning of the NheI-NcoI digested PCR product of the 1863 bp long promoter region of *hmbA* (amplified by “hmbA prom NheI frw” and “hmbA prom NcoI rev” primers). The obtained *NHP6A* expressing vector (pAN-HZS-19) (electronic supplementary material, figure S3*b*) was used for transformation of *hmbAΔ* strain (HZS.320).

In order to obtain *hmbA-gfp* expressing strain (C’*hmbA-gfp*), the *hmbA* gene without stop codon and with an 8 amino acids coding linker sequence at the 3’end (LIDTVDLD) was amplified from wild-type *A. nidulans* genome (HZS.145) by “hmbA NcoI frw” and “hmbA linker NcoI rev” primers and cloned into NcoI site of pAN-HZS-1 vector (Karacsony et al., 2014). The resulted vector (pAN-HZS-20C) (electronic supplementary material, figure S3*b*) was used as template to amplify the 4797 bp long *hmbA-gfp* gene phusion followed by the 3’ UTR of the *trpC* gene (T*_trpC_*) and the *pantoB* selection marker gene by using the chimeric primer pair “hmbA upst chim frw2” and “pantoB hmbA down chim rev”. The *hmbA* –linker – *gfp* - T*_trpC_* - *pantoB* carrying PCR product was joined to the PCR amplified 3428 bp long upstream (“hmbA upst frw”-“hmbA upst rev”) and 3058 bp long downstream (“hmbA down frw”-„hmbA down rev”) regions of *hmbA* by using DJ-PCR method with nested primer pair (“hmbA upst nest frw”-“hmbA down nest rev”) and wild-type *A. nidulans* genome (HZS.145) as template. The assembled PCR product of *hmbA-gfp* cassette was used to transform the *hmbAΔ* strain (HZS.320).

In order to obtain *chiA* overexpressing *hmbAΔ* strain (*hmbAΔ* OE*chiA*), the 2935 bp long *chiA* gene was amplified from wild-type *A. nidulans* genome (HZS.145) by “chiA NcoI fw” and “chiA NotI rev” primer pair. The NcoI-NotI digested PCR product was cloned into NcoI-NotI digested pAN-HZS-1 vector (Karacsony et al., 2014) (the digestion eliminated the *gfp* gene from the vector). The resulted vector (pAN-HZS-31) was used to transform the *hmbAΔ* strain (HZS.320) (electronic supplementary material, figure S3*b*).

After transformation of *hmbAΔ* (HZS.320) with pAN-HZS-9, pAN-HZS-19, the *hmbA-gfp* cassette and pAN-HZS-31, pantothenic acid prototroph transformant strains were collected and the copy number of integration events was measured by qPCR using *hmbA*, *NHP6A*, *gfp*, *chiA* and *pantoB* specific primer pairs (“hmbA ReTi frw”-“hmbA ReTi rev”, “NHP6A ReTi frw”-“NHP6A ReTi rev”, “gfp ReTi frw”-“gfp ReTi rev”, “chiA ReTi frw”-“chiA ReTi rev” and “pantoB ReTi frw”-“pantoB ReTi rev”, respectively). The γ-actin coding *actA* (AN6542) was used as reference single copy gene (amplified by “actA ReTi frw2”-“actA ReTi rev2” primers). The copy number was calculated as described by Herrera et al. (Herrera et al., 2009). The verified transformants C’*hmbA-gfp* (HZS.371, with two copies of transgene), C’*NHP6A* (HZS.834, with one copy of transgene) and *hmbAΔ* OE*chiA* (HZS.921, with one copy of transgene) were used further in the experiments. Complete genotypes of the strains and all the used primers are listed in electronic supplementary material table S1 and S2, respectively.

### Microscopy

For the study of HmbA localization, 10^4^ conidiospores of as C’*hmbA-gfp* (HZS.371) was germinated for 5.5 hours at 37 °C on the surface of cover slips submerged in liquid MM (GLU-NIT). Nuclei were stained with DAPI (4′,6-diamidino-2-phenylindole) (Sigma) according to May (May, 1989). Germlings were studied in Zeiss Axiolab A fluorescent microscope using, Zeiss filter set 15 (GFP) and 49 (DAPI). Chitin staining was carried out by incubating young mycelia grown over cellophane (Ferenczy et al., 1975; Kevei and Peberdy, 1977) in 10 µg/ml Calcofluor White (CFW) stain (Fluka) dissolved in PBS (phosphate buffered saline: 137 mM NaCl, 2.7 mM KCl, 10 mM Na_2_HPO_4_, 1.8 mM KH_2_PO_4_) for 10 min followed by 3 × wash in PBS and microscopy with Zeiss Axioobserver 7 Axiocam 503 mono detector with DAPI filter setting.

### DNA manipulation

Total DNA was prepared from *A. nidulans* as described by Specht et al. (Specht et al., 1982). Transformations of *A. nidulans* protoplasts were performed as described by Antal et al. (Antal et al., 1997). The protoplasts were prepared from mycelia grown over cellophane (Ferenczy et al., 1975; Kevei and Peberdy, 1977) using a 4% solution of Glucanex (Novozymes, Switzerland) in 0.7 M KCl solution. Transformation of 5 × 10^7^ protoplasts was carried out with 100 ng – 1 µg of fusion PCR products. Transformation of *S. cerevisiae* was carried out with the lithium acetate method using single-stranded carrier DNA (Gietz and Schiestl, 2007). Genomic DNA of *S. cerevisiae* was prepared as described by Bodi et al. (Bodi et al., 2017). For cloning procedures, *E. coli* JM109 (Yanisch-Perron et al., 1985) was used and transformation of *E. coli* was performed according to Hanahan (Hanahan, 1983). Plasmid extraction from *E. coli* and other DNA manipulations were done as described by Sambrook et al. (Sambrook, 1989).

### RNA work

For the purpose of Northern analysis, total RNA was isolated from various developmental stages covering the isotropic growth stage of conidiospores (1-3 h incubation), germination and primary hypha formation (from 4 to 5 h), and young mycelia formation (6 h incubation) by a standard protocol using the Trizol Reagent of Invitrogen (CA, USA). Northern blot analysis was performed using the glyoxal method (Sambrook, 1989). Equal RNA loading was calculated by optical density measurements (260/280 nm). [^32^P]-dCTP labelled *hmbA* and *18S rRNA* gene-specific DNA molecules were used as gene probes using the random hexanucleotide-primer kit following the supplier’s instructions (Promega). As a loading control, the expression of *18S rRNA* gene was monitored. For the purpose of mRNA analysis by reverse transcription qPCR (RT-qPCR), total RNA was isolated from yeast from an exponentially growing yeast culture (OD_600_=0.3-1) using RiboPure-Yeast (Invitrogen) kit and from *A. nidulans* mycelia after 9 h incubation using a NucleoSpin RNA Plant kit (Macherey-Nagel) and RNase-Free DNase (Qiagen) according to the manufacturers’ instructions. RNA quality was assessed by using agarose gel electrophoresis. DNA contamination of the RNA samples was checked by performing qPCR on 1 µg RNA samples with *A. nidulans actA* or *S. cerevisiae UBC6* specific primers (“actA ReTi frw2” and “actA ReTi rev2”, “UBC6 ReTi frw” and “UBC6 ReTi rev”) that do not span intron sequences. Samples showing higher than 32 cycle Cq values in the DNA contamination test were used for reverse transcription. cDNA synthesis was carried out with a mixture of oligo-dT and random primers using a RevertAid First Strand cDNA Synthesis kit (Fermentas). RT-qPCR was carried out in a CFX96 Real-Time PCR System (Bio-Rad) with Maxima SYBR Green/Fluorescein qPCR Master Mix (Fermentas) reaction mixture (94 °C for 3 min followed by 40 cycles of 94 °C for 15 s and 60 °C for 1 min). The data processing was performed by the standard curve method (Larionov et al., 2005). The gene expression values of genes of interest were normalized to that of *actA* (γ-actin coding gene AN6542) or *UBC6* (ubiquitin-conjugating enzyme coding gene) reference genes. All the used primers are listed in electronic supplementary material, table S2.

### Extraction and detection of sterigmatocystin (STC) by Thin Layer Chromatography (TLC) and HPLC

Five agar blocks were excised from the centre of 6-day-old *A. nidulans* colonies with a cork borer with 10 mm diameter. STC was extracted from the agar blocks by using 10 ml chloroform. The extracts were concentrated into 1 ml final volume by heating the samples to 65 °C. Samples were loaded on Kieselgel 60 (Merck) plates after being normalized for the protein content of the samples (by using Bradford reagent) and the chromatogram was developed in toluol : ethylacetate : formic acid (50:40:10 V/V/V). Secondary metabolites were detected and recorded under UV light (366 nm), after being sprayed with 10% AlCl_3_ in ethanol, and heated to 100 °C for 1 min. Identification of STC was accomplished by using an STC standard (Sigma).

HPLC detection of STC was carried out using a Shimadzu HPLC system (Shimadzu, Kyoto, Japan) equipped with an SPD-10Avp UV-VIS detector, an LC-20AD binary pump, a SIL-20A autosampler, a DGU-14A degasser, a CTO-10ASvp column thermostat, and a CBM-20A system controller. For data acquisition and evaluation Class VP ver. 6.2 software was used. The separation of sterigmatocystin was performed on a Purosphere Star RP18e, 250 × 4mm, 5 µm column (Merck KGaA, Darmstadt, Germany) at a column temperature of 40 °C. The injected sample volume was 5 µl with a flow rate of 0.5 ml/min. Mixture of water (component A) and methanol (component B) were used as eluents with a gradient program starting with an isocratic step at 60% component B for 1 minute, then increasing to 80% component B in 8 minutes followed by another isocratic step for 16 minutes at 80% component B. After reaching the initial solvent composition in 1 minute, it was held for re-equilibration for 11 minutes. The peak of sterigmatocystin was detected at λ = 254 nm.

### Extraction and detection of trehalose and glycerol by TLC and HPLC

For the monitoring of metabolism of conidial trehalose to glycerol during the swelling of conidiospores that precedes the germ tube formation, 10^9^ conidiospores were inoculated into 200 ml of minimal medium and cultivated at 37 °C with 180 rpm shaking. 40 ml samples were taken after 0, 30, 60, 90 and 120 min of incubation. The conidia were collected by centrifugation (13,000 g, 10 min) and dissolved in 1 ml of 5% (m/V) trichloroacetic acid (TCA). Spores were counted by the use of hemocytometer for the purpose of the normalization of measured trehalose/glycerol content. Sugars and polyalcohols were extracted two times with 1 ml of 5% (m/V) TCA at 80 °C. The solid fractions were removed by centrifugation (13,000 g, 10 min) and the liquid fractions were pooled and concentrated under vacuum to the final volume of 500 μl.

The samples were analysed in separated chromatographic runs for both glycerol and trehalose contents by using the Shimadzu HPLC system as described above (Shimadzu, Kyoto, Japan) except that the original detector was replaced with a refractive index detector (RID-10). In the case of glycerol, the stationary phase was a Hi-Plex H column (300 x 7.7 mm; Agilent, USA) and the mobile phase was 0.005 M H_2_SO_4_ solution at a flow rate of 0.4 ml/min. During the trehalose analysis a silica Si 100 column was used (250 × 4.6 mm; Serva Feinbiochemica, Germany) and the components were eluted with 17/83 V/V mixture of water and acetonitrile at the flow rate of 1.2 ml/min. In both cases, the injection volume was 20 µl and the cell of the detector was tempered at 55 °C. The run time of the glycerol and trehalose HPLC methods was 30 min and 15 min, while the column temperature was maintained at 60 °C and 35 °C, respectively. The retention time of glycerol and trehalose peaks in the two separated method were 19.9 min and 7.3 min, respectively.

### Protein modelling

Initial protein models were obtained by I-Tasser (Zhang, 2008), whereas the refined models were obtained by ModRefiner (Xu and Zhang, 2011). The obtained pdb files were further analysed in UCSF Chimera 1.14. Ramachandran plot analysis was carried out by using Procheck server (https://servicesn.mbi.ucla.edu/PROCHECK/). Electrostatic potential was calculated with Delphi Web Server with the default parameters (http://compbio.clemson.edu/sapp/delphi_webserver/) (Sarkar et al., 2013; Smith et al., 2012). The resulted pqr file was submitted to Chimera and electrostatic surface colouring was done by the Coulombic Surface Colouring application of Chimera.

### *S. cerevisiae* fitness measurements

Growth was assayed by monitoring the optical density at 600 nm (OD_600_ value) of liquid cultures using 384 well density microtiter plates following a previous protocol (Kovacs et al., 2021). The growth curves were monitored over a 72 h incubation period in a Powerwave HT plate reader (BioTek Instruments Inc). During the kinetic run, the optical density of each well was recorded at 600 nm (OD_600_) every 5 minutes. Between the optical readings, the cultures were incubated at 30 °C, with alternating shaking speed (1000-1200 rpm). A modified version of a published procedure (Warringer and Blomberg, 2003; Warringer et al., 2003), implemented in R (R Core Team, 2021) was used to estimate several fitness components, including growth rate, doubling time, optical density increment, and length of the lag phase.

## Electronic supplementary material

### Supplementary tables

**Supplementary table S1.** List of *A. nidulans* and *S. cerevisiae* strains used in this work.

**Supplementary table S2.** List of primers used in this study.

### Supplementary figures

**Supplementary Figure S1.** Response of *hmbAΔ* and the reconstituted strains to various environmental conditions.

**Supplementary Figure S2.** Presentation of micromorphology of hypha-elongation in *hmbA^+^* control, *hmbAΔ* and in the various reconstituted strains.

**Supplementary Figure S3.** Schematic presentation of the vectors constructed in this work.

### Supplementary data

**Supplementary data.** Colony size (mm) and the calculated relative growth of *hmbA^+^*, *hmbAΔ* and complemented control strains (C’*hmbA-gfp*, C’*hmbA* and C’*NHP6A*) on various carbon-sources, nitrogen-sources, on various temperatures, osmotic stabilizer and in the presence of osmotic-, oxidative-, cell-wall- and heavy metal stressors.

## Supporting information

electronic supplementary material, figures S1-S3

electronic supplementary material, tables S1, S2

## Acknowledgement

The project has received funding from the EU’s Horizon 2020 research and innovation program under grant agreement No. 739593. Work was supported by the Hungarian National Research, Development and Innovation Office (NKFIH K16-119516) and by the Hungarian Government (GINOP-2.3.2-15-2016-00012).

