## Supplementary material for "Role of High Mobility Group B protein HmbA, orthologue of yeast Nhp6p in *Aspergillus nidulans*": electronic supplementary material, figures S1-S3

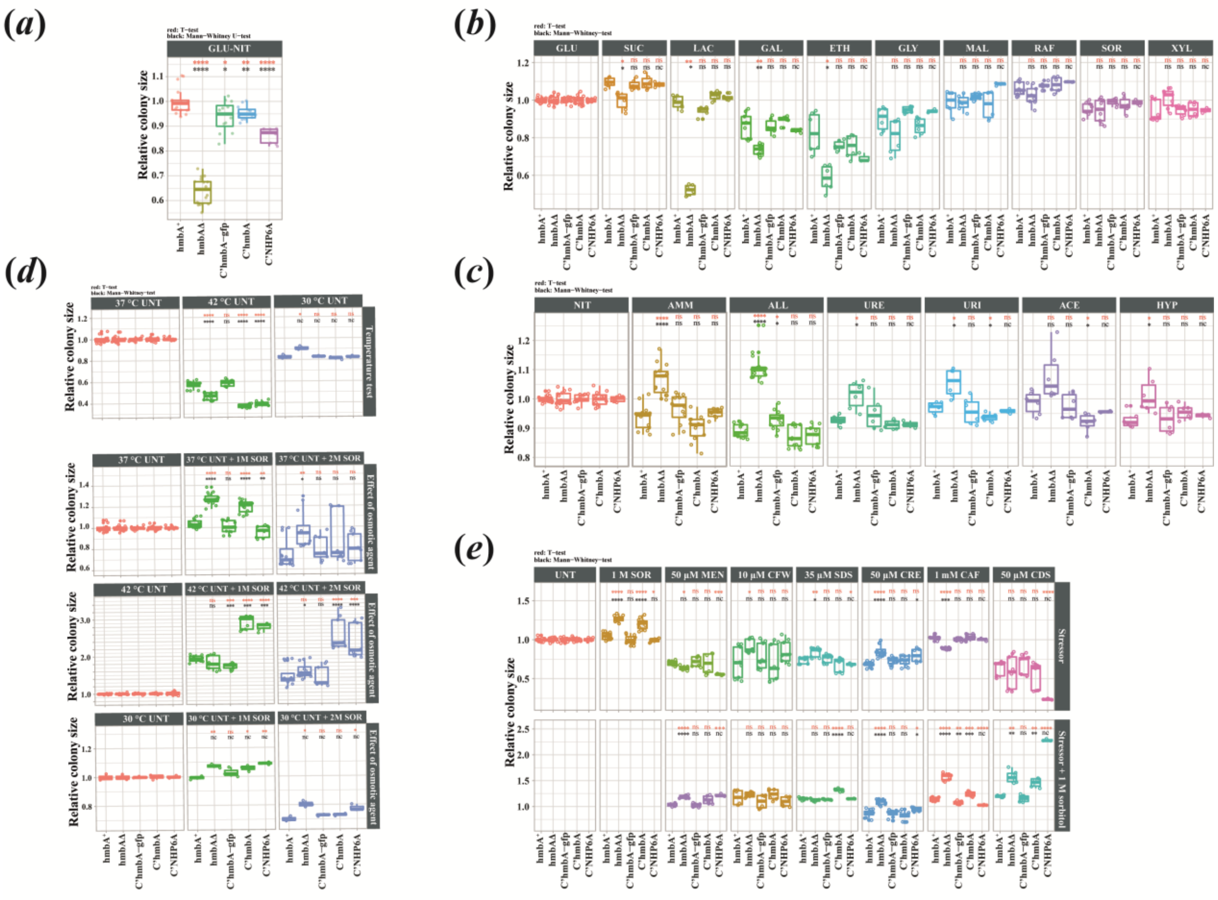


**Supplementary Figure S1.** Response of *hmbAΔ* and the reconstituted strains to various environmental conditions. Response of all examined strains to the tested conditions was calculated by strainwise normalization of the colony sizes to the control condition. The calculated relative colony size were implemented in R (R Core Team, 2021). (*a*) Comparison of growth of *hmbAΔ* and the reconstituted strains to the growth of *hmbA*$^+$ control strain on glucose-sodium nitrate (GLU-NIT) minimal medium. To assess significant differences between the growth of *hmbA*$^+$ control and the tested strains, we used Student's *t*-test (red asterisks) and Mann-Whitney U-test (black asterisks): * $P < 0.05$; ** $P < 0.01$; **** $P < 0.0001$. (*b*) Study of the utilization ability of various carbon-sources. In all minimal medium, the nitrogen-source was sodium nitrate. The control condition was the GLU (glucose). To assess significant differences in the response of *hmbAΔ* and in the various reconstituted strains to that of the *hmbA*$^+$ control we used Student's *t*-test (red asterisks/letters) and Mann-Whitney U-test (black asterisks/letters): * $P < 0.05$; ** $P < 0.01$; nc: non-calculated due to low sample size; ns: non-significant. SUC: sucrose; LAC: lactose; GAL: galactose and ETH: ethanol; GLY: glycerol, MAL: maltose; RAF: raffinose; SOR: sorbitol and XYL: xylose. (*c*) Study of the utilization ability of various nitrogen-sources. In all minimal medium, the carbon-source was glucose. The control condition was the NIT (sodium nitrate). To assess significant differences in the response of *hmbAΔ* and in the various reconstituted strains to that of the *hmbA*$^+$ control we used Student's *t*-test (red asterisks/letters) and Mann-Whitney U-test (black asterisks/letters): * $P < 0.05$; **** $P < 0.0001$; nc: non-calculated due to low sample size; ns: non-significant. AMM: diammonium L-(+)-tartrate; ALL: allantoin; URE: urea; URI: uric acid; ACE: acetamide and HYP: hypoxanthine. (*d*) Study of the effect of various temperatures, osmotic stabilizer (1 M sorbitol) and osmotic stressor (2 M sorbitol) to the growth ability. All minimal medium was GLU-NIT. The control condition was the condition indicated above the first column in each box plot. They were 37 °C UNT in the first and second box plot (37 °C untreated condition), 42 °C UNT in the third box plot (42 °C untreated condition) and 30 °C UNT in the fourth box plot (30 °C untreated condition). To assess significant differences in the response of *hmbAΔ* and in the various reconstituted strains to that of the $hmbA^+$ control we used Student's *t*-test (red asterisks/letters) and Mann-Whitney U-test (black asterisks/letters): * $P < 0.05$; ** $P < 0.01$; *** $P < 0.001$; **** $P < 0.0001$; nc: non-calculated due to low sample size; ns: non-significant. (*e*) Study of the effect of the supplementation of the medium with various stressors on the growth ability without (first row of box plots) and with (second row of box plots) osmotic stabilizer (1 M SOR (sorbitol)). All minimal medium was GLU-NIT. The control condition was the untreated GLU-NIT medium for the first row of box plot, while the control condition for the second row of box plots was GLU-NIT medium supplemented with 1 M sorbitol (1 M SOR). To assess significant differences in the response of *hmbAΔ* and in the various reconstituted strains to that of the $hmbA^+$ control we used Student's *t*-test (red asterisks/letters) and Mann-Whitney U-test (black asterisks/letters): * $P < 0.05$; ** $P < 0.01$; *** $P < 0.001$; **** $P < 0.0001$; nc: non-calculated due to low sample size; ns: non-significant. MEN: menadione; CFW: Calcofluor White; SDS: sodium dodecyl sulphate; CRE: congo red; CAF: caffeine and CDS: cadmium sulphate. Concentrations of the stressors are indicated above the box plots. Box plots show the median, first and third quartiles, with whiskers showing the $5^{th}$ and $95^{th}$ percentiles. The experiments were executed at least in three biological replicates. All raw data is available in electronic supplementary material, data file. Concentrations of used sole carbon- and nitrogen-sources are listed in the Materials and methods section of the main text. Used strains were $hmbA^+$ as control (HZS.120); *hmbAΔ* (*hmbA* deletion strain, HZS.320); *C'hmbA* (*hmbAΔ* complemented with *hmbA*, HZS.621), C'*hmbA-gfp* (*hmbAΔ* complemented with *hmbA-gfp*, HZS.371) and C'*NHP6A* (*hmbAΔ* complemented with *NHP6A*, HZS.834). Complete genotypes are listed in the electronic supplementary material, table S1.

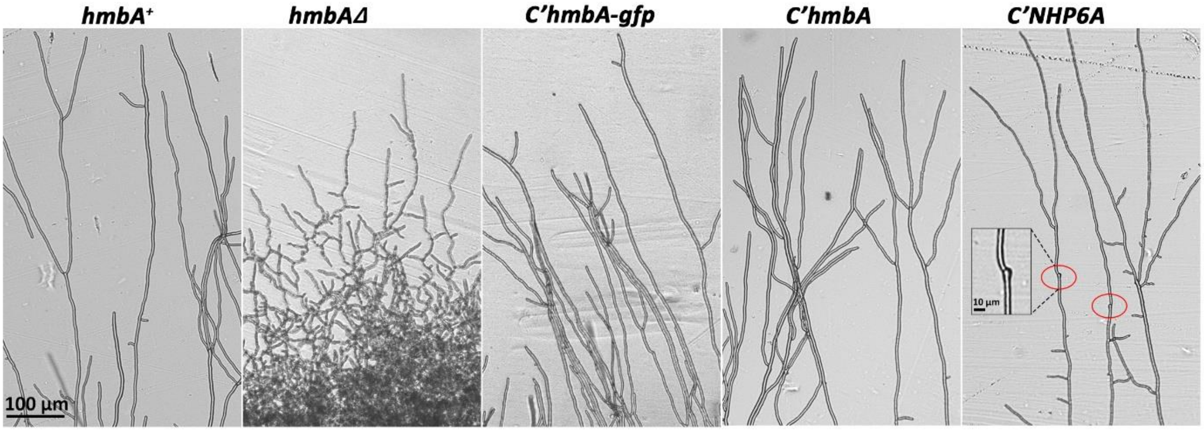


**Supplementary Figure S2.** Presentation of micromorphology of hypha-elongation in *hmbA*$^{+}$ control, *hmbAΔ* and in the various reconstituted strains. Brightfield images were documented with Leica DMI 4000B, DFC295 detector. Scale bar is shown. Strains were grown on glucose-nitrate minimal medium at 37 °C for 30 h. Red circles: mark the relicts of arrested hyphal tips in C'*NHP6A* strain that underwent a germination-like process to continue the polar growth (this phenotype is typical for *hmbAΔ* strain. Strains were *hmbA*$^{+}$ as control (HZS.120); *hmbAΔ* (*hmbA* deletion strain, HZS.320); *C'hmbA* (*hmbAΔ* complemented with *hmbA*, HZS.621), C'*hmbA-gfp* (*hmbAΔ* complemented with *hmbA-gfp*, HZS.371); C'*NHP6A* (*hmbAΔ* complemented with *NHP6A*, HZS.834).

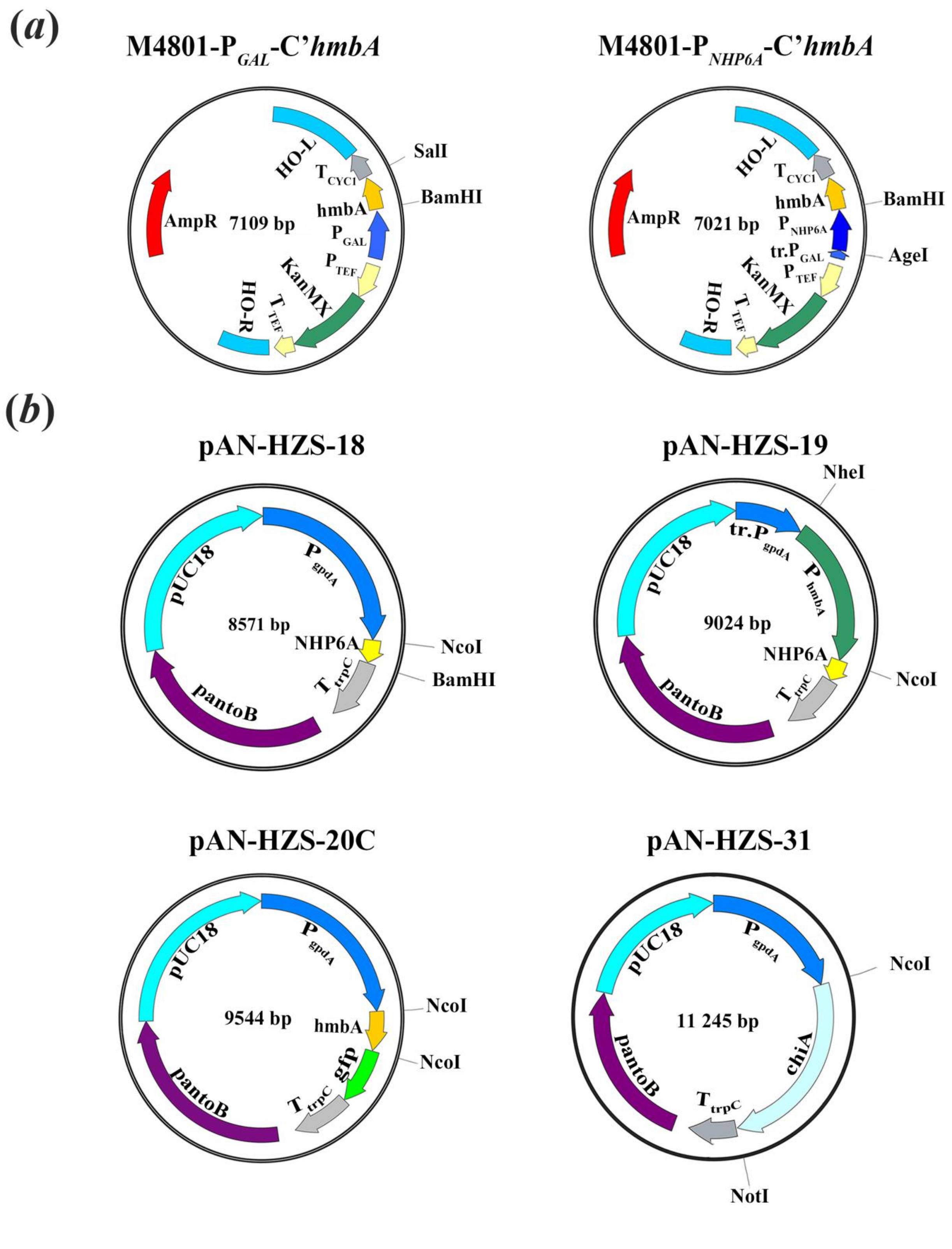


**Supplementary Figure S3.** Schematic presentation of the vectors constructed in this work. (*a*) Schematic presentation of the yeast transformation vectors developed from the modification of M4801 vector (http://www.addgene.org/51664/). For the detailed description of the construction process, see the Materials and methods section of the main text. Names of the vectors are shown above the circular schemes, the size of the vectors are shown within the circular schemes. Restriction sites used for cloning are shown. Coloured arrows show relevant components of the vectors, the arrowheads indicate the orientation. HO-L and HO-R: left (L) and right (R) arm of the *HO* gene that encodes a DNA endonuclease responsible for mating-type switch via the formation of a double-strand break at the mating-type locus*;* $T_{CYC1}$: termination sequence of *CYC1* (Cytochrome C1); hmbA: coding gene of HmbA from *A. nidulans*; $P_{GAL}$: galactose-inducible promoter; $P_{TEF}$ and $T_{TEF}$: promoter and termination sequence of *TEF* gene (Translational elongation factor EF-1) from *S. cerevisiae;* KanMX: kanamycin resistance marker gene; AmpR: ampicillin resistance marker gene; tr.$P_{GAL}$:truncated *GAL* promoter; $P_{NHP6A}$: native *NHP6A* promoter from *S. cerevisiae*. M4801-$P_{NHP6A}$-C'*hmbA* was used for transformation of HZS.891 and the obtained yeast reconstitution strain was named as yC'*hmbA* (HZS.890). (*b*) Schematic presentation of the *A. nidulans* transformation vectors developed from pAN-HZS-1 (Karacsony et al., 2014). For the detailed description of the construction process, see the Materials and methods section of the main text. pAN-HZS-18 vector was used for the construction of the transformation vector pAN-HZS-19. pAN-HZS-20C was used as template for the construction of the transformation cassette *hmbA-gfp* obtained by using double-joint PCR method (see Materials and methods section of the main text). Unique restriction motifs used for cloning are shown. Coloured arrows show relevant components of the vectors, the arrowheads indicate the orientation. pUC18: standard *E. coli* vector; $P_{gpdA}$: constitutive promoter of *gpdA* (glyceraldehyde-3-phosphate dehydrogenase coding gene) from *A. nidulans*; gfp: coding gene of Gfp (green fluorescence protein); $T_{trpC}$: termination sequence of *trpC* gene (tryptophan biosynthesis gene) from *A. nidulans*; pantoB: wild-type *pantoB* gene (coding for pantothenic acid biosynthesis gene) from *A. nidulans*, which serves as selection marker gene for transformation; tr.$P_{gpdA}$: truncated *gpdA* promoter; hmbA*:* coding gene of HmbA from *A. nidulans;* $P_{hmbA}$: physiological promoter of *hmbA*;. NHP6A: coding gene of Nhp6Ap from *S. cerevisiae*; chiA: coding gene of ChiA from *A. nidulans.* Names of the vectors are shown above the circular schemes, the size of the vectors are shown within the circular schemes. These vectors were used for transformations of *A. nidulans* recipient strains (see Materials and methods section of the main text) to obtain the C'*NHP6A* (HZS.834) and C'*hmbA-gfp* (HZS.371) *A. nidulans* reconstitution strains and the *chiA* overexpression *hmbA*Δ OE*chiA* strain (HZS.834).

R Core Team, R: A language and environment for statistical computing. In: V. Foundation for Statistical Computing, Austria, (Ed.), 2021.
