## Supplementary material for "Role of High Mobility Group B protein HmbA, orthologue of yeast Nhp6p in *Aspergillus nidulans*": electronic supplementary material, tables S1, S2

**Supplementary Table S1. List of *A. nidulans* and *S. cerevisiae* strains used in this work.**

All *Aspergillus* strains listed are *veA1* mutant.

| ***A. nidulans* strains** | | | |
|---|---|---|---|
| Strain | Genotype | Purpose | Reference |
| HZS.120 | *riboB2 pabaA1* | growth test; microscopic experiments; mRNA expression analysis; metabolite analysis | [1] |
| HZS.145 | *veA1* | template DNA for PCRs | [2] |
| HZS.320 | *hmbAΔ::riboB*$^{+}$ *pantoB100 pabaA1 riboB2* | recipient strain for transformation experiment to obtain C'*NHP6A,* C'*hmbA-gfp* and *hmbAΔ* OE*chiA* strains; growth test; microscopic experiments; mRNA expression analysis; metabolite analysis | [3] |
| HZS.371 | *hmbAΔ::riboB*$^{+}$ *pantoB100 pabaA1 riboB2* +*in trans* "*hmbA-gfp* cassette" in 2 copy | microscopic experiments; growth test; metabolite analysis | this work (by transformation of the "*hmbA -gfp* cassette" into HZS.320) |
| HZS.621 | *hmbAΔ::riboB*$^{+}$ *pantoB100 pabaA1 riboB2* +pAN-HZS-9 plasmid integrated into the *hmbA* promoter in 1 copy | growth test; metabolite analysis | [3] |
| HZS.834 | *hmbAΔ::riboB*$^{+}$ *pantoB100 pabaA1 riboB2* + *in trans* pAN-HZS-19 plasmid in 1 copy | growth test; metabolite analysis | this work (by transformation of the pAN-HZS-19 plasmid into HZS.320) |
| HZS.921 | *hmbAΔ::riboB*$^{+}$ *pantoB100 pabaA1 riboB2 +in trans* pAN-HZS-31 | microscopic experiments; metabolite analysis | this work (by transformation of the pAN-HZS-31 plasmid into HZS.320) |
| ***S. cerevisiae* strains** | | | |
| Strain | Genotype | Purpose | Reference |
| NHP6AΔ | *nhp6aΔ::KanMX his3Δ1 leu2Δ0 met15Δ0 ura3Δ0 MAT-a* | recipient strain for transformation experiment to obtain NHP6AΔ (HphMX) strain | [4] |

| | | | |
|---|---|---|---|
| NHP6BΔ | *nhp6bΔ::NatMX can1Δ::p$_{STE2}$-SpHis5 lyp1Δ his3Δ1 leu2Δ0 ura3Δ0 met15Δ0 MAT- α* | parental strain in genetic crosses with NHP6AΔ (HphMX) | [5] |
| NHP6AΔ (HphMX) | *nhp6aΔ::HphMX his3Δ1 leu2Δ0 met15Δ0 ura3Δ0 MAT-a* | parental strain in genetic crosses with NHP6BΔ | this work (by transformation of the pCRII-TOPO::hphMX plasmid into NHP6AΔ strain) |
| HZS.890 | *nhp6aΔ::HphMX nhp6bΔ::NatMX his3Δ1 leu2Δ0 met15Δ0 ura3Δ0 +in trans* M4801-P$_{NHP6A}$-C'*hmbA* plasmid in 1 copy | mRNA expression analysis; growth test; fitness measurements | this work (by transformation of the M4801-P$_{NHP6A}$-C'*hmbA* plasmid into HZS.891) |
| HZS.891 | *nhp6aΔ::HphMX nhp6bΔ::NatMX his3Δ1 leu2Δ0 met15Δ0 ura3Δ0* | recipient strain for transformation experiment to obtain yC'*hmbA* strain; mRNA expression analysis; growth test; fitness measurements | this work (obtained by genetic cross of NHP6AΔ (HphMX) and NHP6BΔ) |
| Y199 | *his3Δ::KanMX hoΔ::NatMX his3Δ1 leu2Δ0 met15Δ0 ura3Δ0 MAT a* | mRNA expression analysis; growth test; fitness measurements | provided by Zoltán Farkas |

Explanation of mutant alleles, which are not described in the text: *veA1* is a mutation in the *veA* gene resulting profuse conidiation regardless of the presence or absence of light [6]. $T_{trpc}$: terminator sequence of trpC gene. *KanMX* (G418), *NatMX* (nourseothricin) and *HphMX* (hygromycin B) symbols are resistance marker genes. *lyp1Δ:* deletion of a lysine permease responsible for uptake of cationic amino acids. *hoΔ:* deletion of HO (YDL227C) made by allelic replacement using the *ho::KanMX* deletion cassette. *HO* is a gene encoding a DNA endonuclease responsible for mating-type switch *via* the formation of a double-strand break at the mating-type locus. *can1Δ*: deletion of *CAN1* (YEL063C) made by allelic replacement using the *can1::p$_{STE2}$-SpHis5* deletion cassette. *CAN1* is a gene encoding arginine permease and its deletion confers resistance to the toxic arginine analog, canavanine. The selectable marker, in this case, is the p$_{STE2}$-*SpHis5*, which is the His5p of *Schizosaccharomyces pombe* under the control of the promoter of *STE2* (p$_{STE2}$), that is active only in matA *S. cerevisiae* cells. Other gene symbols refer to auxotrophies: *pabaA1*, p-aminobenzoic acid; *pantoB100*, pantothenic acid and *riboB2*, riboflavin; *his3Δ1*: histidine; *leu2Δ0*: leucine; *met15Δ0*: methionine; *ura3Δ0*: uracil, uridin.

**Supplementary Table S2. List of primers used in this study.**

| | |
|---|---|
| **C'*NHP6A* strain** | |
| nhp6A NcoI frw | 5'-ttttttttccatggtcaccccaagagaacctaag-3' |
| nhp6A BamHI rev | 5'-ttttttttggatccctaagccaaagtggcgttatataactc-3' |
| hmbA prom NheI frw | 5'-ttttttttgctagcgatcctcaatgaaccttgcccttg-3' |
| hmbA prom NcoI rev | 5'-ttttttttccatggggtgaaggtctgaagctgttggacg-3' |
| **C'*hmbA-gfp* strain** | |
| hmbA NcoI frw | 5'-ttttttttccatggctaaggccaatcctac-3' |
| hmbA linker NcoI rev | 5'-ttttttttccatggaatcaagatcgactgtatcaataagggacgactcctcatcctcttcg-3' |
| hmbA upst chim frw2 | 5'-*cttcgtccaacagcttcagaccttcacca*tggctaaggccaatcctacccg-3' |
| pantoB hmbA down chim rev | 5'-*cggaggtcaagcgactcgacactaaggtg*acataatcttatgatccataccacctagc-3' |
| hmbA upst frw | 5'-ctctgacctgccacgaggccttgctctatg-3' |
| hmbA upst rev | 5'-ggtgaaggtctgaagctgttggacgaagag-3' |
| hmbA down frw | 5'-caccttagtgtcgagtcgcttg-3' |
| hmbA down rev | 5'-gcatacaatgcgagcacggtggtcgtcatc-3' |
| hmbA upst nest frw | 5'-ggagacatttcgaactgtatcagggctaac-3' |
| hmbA down nest rev | 5'-cggtgctgttgctgcttgaggacgaggag-3' |
| ***hmbAΔ* OE*chiA*** | |
| chiA NcoI fw | 5'-ttttttttccatggcccctaaactgtttaccttc-3' |
| chiA NotI rev | 5'-ttttttttgcggccgcttacaaaacagcaagcagggagag-3' |
| **yC'*hmbA*** | |
| c-hmbA BamHI frw | 5'-ttttttttggatccatgcctaaggccaatcctacc-3' |
| c-hmbA SalI rev | 5'-ttttttttgtcgacttaggacgactcctcatcctcttc-3' |
| NHP6A prom AgeI frw | 5'-ttttttttaccggttcttggagcgttgagcacgtctac-3' |
| NHP6A prom BamHI rev | 5'-ttttttttggatcctgcgactgtgctttactatgtatagggtag-3' |
| **Checking gene deletions and integrations** | |
| NHP6A confA frw | 5'-cacgacgttaaatactgttcaagtg-3' |
| NHP6B confB frw | 5'-ccacctctcacccaatattctg-3' |
| KanB rev | 5'-ctgcagcgaggagccgtaat-3' |
| NHP6A prom AgeI frw | 5'-ttttttttaccggttcttggagcgttgagcacgtctac-3' |
| NHP6A prom BamHI rev | 5'-ttttttttggatcctgcgactgtgctttactatgtatagggtag-3' |
| **qPCR** | |
| hmbA ReTi frw | 5'-aaagatgctcggtgagaagtg-3' |
| hmbA ReTi rev | 5'-ctcgtaccgcttcttgtcag-3' |
| NHP6A ReTi frw | 5'-agaagttgggtgagaagtgg-3' |
| NHP6A ReTi rev | 5'-ttcatatctcttcttatcggcctg-3' |
| gfp ReTi frw | 5'-atcttcttcaaggacgacgg-3' |
| gfp ReTi rev | 5'-ttgaagtcgatgcccttcag-3' |
| chiA ReTi frw | 5'-ctcttcaagcacttccactc-3' |
| chiA ReTi rev | 5'-gccagatgatgtacttgtagag-3' |
| pantoB ReTi frw | 5'-gttaagagccgagcgtatcc-3' |
| pantoB ReTi rev | 5'-cttcaggtaattcatcaacagcc-3' |
| actA ReTi frw2 | 5'-accatgtaccctggtatctc-3' |
| actA ReTi rev2 | 5'-ggaggagcaatgatcttgac-3' |
| UBC6 ReTi frw | 5'-ttacaagggcggtcaatatcac-3' |
| UBC6 ReTi rev | 5'-gggtggtaatcactcatagaaagg-3' |

| **RT-qPCR** | |
|---|---|
| actA ReTi frw | 5'-ggtatcatgatcggtatggg-3' |
| actA ReTi rev | 5'-tatctgagtgtgaggatacca-3' |
| UBC6 ReTi frw | 5'-ttacaagggcggtcaatatcac-3' |
| UBC6 ReTi rev | 5'-gggtggtaatcactcatagaaagg-3' |
| chiA ReTi rev | 5'-gccagatgatgtacttgtagag-3' |
| chiA ReTi frw | 5'-ctcttcaagcacttccactc-3' |
| SNR6 ReTi frw | 5'-cgaagtaacccttcgtggac-3' |
| SNR6 ReTi rev | 5'-aacggttcatccttatgcagg-3' |

Underlined letters in the primer sequences refer to the restriction sites designed within.
Italic letters at the 5' end refer to the chimeric nature of the primer.
